# Characterization of proprioceptive system dynamics in behaving *Drosophila* larvae using high-speed volumetric microscopy

**DOI:** 10.1101/467274

**Authors:** Rebecca Vaadia, Wenze Li, Venkatakaushik Voleti, Aditi Singhania, Elizabeth M.C. Hillman, Wesley B. Grueber

## Abstract

Proprioceptors provide feedback about body position that is essential for coordinated movement. Proprioceptive sensing of the position of rigid joints has been described in detail in several systems, however it is not known how animals with an elastic skeleton encode their body positions. Understanding how diverse larval body positions are dynamically encoded requires knowledge of proprioceptor activity patterns *in vivo* during natural movement. Here we applied high-speed volumetric SCAPE microscopy to simultaneously track the position, physical deformation, and temporal patterns of intracellular calcium activity of multidendritic proprioceptors in crawling *Drosophila* larvae. During the periodic segment contraction and relaxation that occurs during crawling, proprioceptors with diverse morphologies showed sequential onset of activity throughout each periodic episode. A majority of these proprioceptors showed activity during segment contraction with one neuron type activated by segment extension. Different timing of activity of contraction-sensing proprioceptors was related to distinct dendrite terminal targeting, providing a continuum of position encoding during all phases of crawling. These dynamics could endow different proprioceptors with specific roles in monitoring the progression of contraction waves, as well as body shape during other behaviors. We provide activity measurements during exploration as one example. Our results provide powerful new insights into the body-wide neuronal dynamics of the proprioceptive system in crawling *Drosophila*, and demonstrate the utility of our approach for characterization of neural encoding throughout the nervous system of a freely behaving animal.

## Introduction

Monitoring of neural activity in freely behaving animals is a key step towards understanding how sensory activity is transformed into action [1-3]. Small invertebrate model systems with well-described sensory systems and complete or near-complete connectomes, such as *C. elegans* and *Drosophila* larvae, are ideal systems in which to uncover fundamental principles of sensorimotor integration. Light sheet, confocal and two-photon microscopy can capture neuronal calcium activity in isolated *Drosophila* brains or immobilized preparations [4-8]. However, these methods have been unable to provide volumetric imaging at sufficient speeds, in unrestrained samples, to enable extended imaging of body-wide neural activity in behaving animals.

Here, we applied multi-spectral, high-speed, volumetric Swept Confocally Aligned Planar Excitation (SCAPE) microscopy [9, 10] to characterize the dynamics of neuronal activity in crawling *Drosophila* larvae. We focused on the *Drosophila* multidendritic (md) neurons, which are located just within the larval body wall and innervate connective tissues [11, 12], the epidermis [13, 14] and internal nerves [15]. A subset of six md neurons (Fig. 1a) extend axons to more dorsal neuropil regions important for motor control, suggesting that they are proprioceptors that provide feedback on body position [15-18]. This feedback is thought to be particularly important during crawling, which involves periodic strides driven by segmental muscle contractions progressing from posterior to anterior along the body [19]. However, studies investigating the activity of these sensors have been limited to dissected preparations: imaging of axon terminals in an isolated central nervous system (CNS) suggests that at least some of these neurons are active during muscle contraction [20], while an electrophysiology study has shown activity in one cell type in response to stretch in a dissected preparation [21]. Studies that disabled all or some of these six neurons observed significantly slowed crawling, suggesting that these cells are proprioceptors that provide a segment contraction “mission accomplished” signal that promotes progression of the peristaltic wave [14]. These behavioral studies concluded that this set of neurons have redundant functions during crawling, because silencing different subsets caused similar behavioral deficits. However, the diverse dendrite morphologies and positions of these proprioceptor neurons [13, 15] suggest that each is likely to have distinct sensitivities and functions. Identifying the specific roles of each cell type is not possible without measuring the system’s dynamic activity patterns during natural movements to provide a more comprehensive view of the synergies and dynamic encoding properties of the larval proprioceptive circuit.

**Figure 1.**
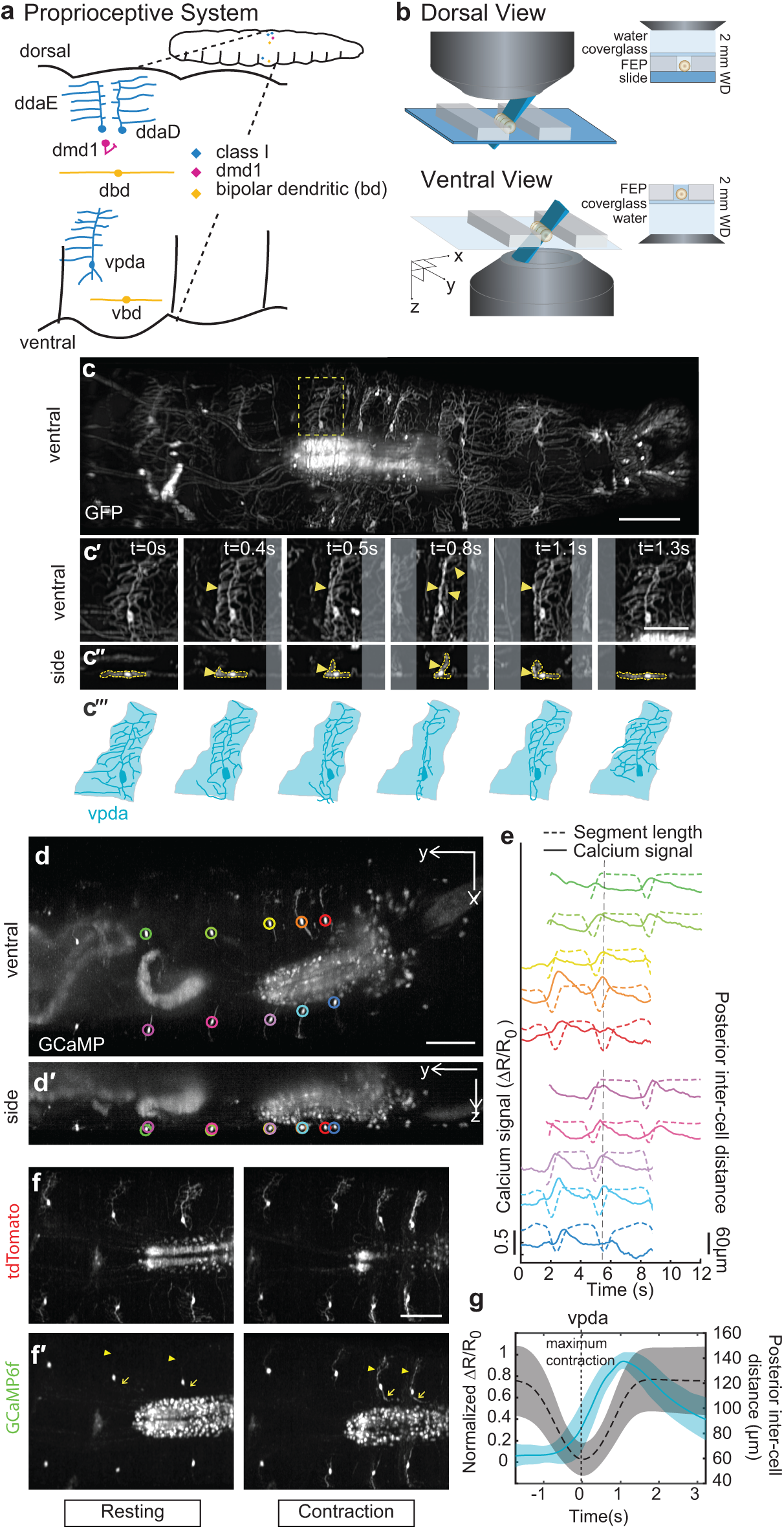
SCAPE imaging of proprioceptor dendrite and activity dynamics in crawling larvae. (**a**) Schematic of the larval proprioceptive system. **(b-b’)** Schematic of larval imaging platform for SCAPE microscopy. An upright configuration is used to image the dorsal side of the larva, and an inverted configuration is used to image the ventral side. (**c**) SCAPE imaging of *221*-*Gal4*, *UAS*-*CD4tdGFP* larva ventral side during crawling, Maximum Intensity Projection (MIP) over a 95 μm depth range from a 160μm deep volume (to exclude gut autofluorescence, square root grayscale – see methods). supplemental movie 1 shows a ventral dataset in full. Yellow box indicates neuron examined in time lapse in **c’-c’’. (c’-c’’)** vpda in ventral view (**c’)** and side view (**c’’)** in successive time lapse frames during forward crawling. vpda dendritic arbors and cell body are outlined in yellow. Other arbors are from cIV neurons. (**c’’’)** Tracing of neurons in c’. Shaded areas represent dendritic field territory at t=0s before folding. **(d-d’)** SCAPE GCaMP imaging of a *410*-*Gal4*, *20XUAS*-*IVS*-*GCaMP6f* (x2), *UAS*-*CD4tdTomato* larva, ventral side, during forward crawling. Images are MIP of a full 168μm deep imaging volume (square root grayscale). **(d)** shows an x-y view and (**d**’) shows side view (y-z). GCaMP signal was extracted from the segmentally repeated vpda neurons indicated by colored circles. **(e)** vpda soma response for the neurons tracked in **(d)**. The distance between the measured neuron and the posterior neuron (posterior inter-cell distance) is plotted in dashed lines and the GCaMP6f response is plotted in solid lines (quantified as the fractional change in the green / red fluorescence ratio ΔR/R_0_ – see methods and see Fig. S1). Dotted vertical line refers to time point shown in **d**. An animated version of this dataset is shown in supplemental movie 2. **(f)** tdTomato and **(f’)** GCaMP6f images showing representative example of increased activity in dendrites (arrowhead) and axons (arrow) during contraction. Images are cropped to show region of interest and are MIP of a 70 μm depth range from a 160 μm deep volume (square root grayscale). See supplemental movie 3. **(g)** Mean calcium response (solid line) ± standard deviation (s.d., ribbon) of vpda soma (3 animals, n=22 cells (8, 10 and 4), 26 events (9, 12 and 4 respectively)) during segment contraction, represented by mean posterior inter-cell distance (dashed line) ± s.d. (ribbon). Maximal posterior segment contraction is set at ‘t=0s’ for each recording. ΔR/R0 amplitude was normalized for each event. Posterior is to the left for all images. Scale bar=100 μm in c, d, f, Scale bars=50 μm in c’, c’’.

Here, we characterized the spatiotemporal and functional dynamics of this set of *Drosophila* md proprioceptors by imaging neurons co-expressing GCaMP and tdTomato using SCAPE microscopy, with subsequent dynamic tracking and ratiometric calcium signal extraction. Characterization of the real-time dynamics of segment contraction and extension during crawling and exploratory head movements revealed that proprioceptors increased their calcium levels in synchrony with deformation of their dendrites. These cells provide a striking sequence of signaling during stereotyped forward crawling, inconsistent with redundancy, but rather suggesting an elegant continuum of sensing during the movement. Furthermore, analysis of sensory responses during non-stereotyped exploration revealed a linear combinatorial code for the representation of complex, simultaneous turning and retracting movements. These results provide valuable new input for models of how movements are controlled via feedback in the context of the larval connectome, and also demonstrate a new approach for characterization of body-wide neuronal dynamics in behaving *Drosophila*.

## Results

### SCAPE microscopy allows 3D imaging of dendritic deformations and activity dynamics in behaving larvae

To begin to characterize proprioceptor dynamics as larvae crawl, we focused on the ventral class I (cI) neuron vpda (Fig. 1a). cI neurons spread sensory dendrites along the body wall epidermis, suggesting that these cells may detect cuticle folding. To investigate how vpda sensory terminals deform during crawling, we first characterized dendrite dynamics using high-speed volumetric SCAPE microscopy [9, 10] at 10 volumes per second (VPS) as a larva crawled within a linear channel (Fig. 1b-c). To achieve this imaging, we made numerous improvements to our original SCAPE microscopy system including substantially improving spatial resolution to permit individual dendrites to be clearly resolved in 3D at high speed in the freely moving larva. We also increased the field of view to over 1mm and made it sufficiently uniform to capture the entire crawling larva (see methods).

During peristalsis, vpda proprioceptors expressing GFP as a static marker showed repeated folding and extension; folding as a peristaltic wave entered a segment, and extending as the wave moved to anterior segments (Fig.1c-c’’’, supplemental movie 1). vpda dendrites viewed in cross section showed hinge-like dynamics, sweeping across an approximately 90° angle during each peristaltic contraction (Fig. 1c”). These data indicate that vpda dendrites are positioned to respond to the repeated folding and extension of the body wall that occurs during crawling.

Next, we sought to reveal if and how the activity of these neurons changes as the dendrites fold during segment contraction. If vpda neurons indeed function as proprioceptors, we should be able to detect activity in these cells during locomotion. We built a dual-expression line of larvae to label targeted proprioceptive cells with both calcium-sensitive GCaMP (green) and static tdTomato (red). To acquire SCAPE microscopy data in this model, we optimized parallel dual color imaging and developed a tracking algorithm that localizes the cell bodies via their static red fluorescence. These tracked cells can then be used as fiducials for quantification of movement and behavior, as well as to extract and ratiometrically correct simultaneously recorded GCaMP fluorescence from the same cells.

We tracked inter-cell distance (defined as the distance between the measured neuron and a homologous neuron in the posterior or anterior segment) to relate calcium dynamics to segmental contraction and extension phases. We observed rises in GCaMP fluorescence in vpda neurons during segment contraction (Fig. 1d-e, g). Calcium signals subsided as the peristaltic wave progressed to adjacent anterior segments (Fig 1d-e, g; supplemental movies 2-3). Dynamic calcium responses were also visible in dendritic arbors and axons (Fig. 1f arrowheads and arrows).

As a control, an additional larva line co-expressing GFP and tdTomato in vpda neurons was imaged using SCAPE during crawling. Applying the same tracking and ratiometric correction as for GCaMP, we observed insignificant changes in ratiometrically corrected GFP signal during crawling (Fig. S1). Taken together, our data indicate that vpda neurons respond to body wall folding during segment contraction.

### Monitoring of different proprioceptive cell types reveals distinct activity patterns

Having established this pipeline for cell characterization, SCAPE was then used to monitor the physical and functional dynamics of the remaining proprioceptive cell types, each of which has unique dendrite morphologies and positions (Fig. 1a). Two additional cI neurons besides vpda project secondary dendrites along the dorsal side of the body wall (ddaD anteriorly and ddaE posteriorly; Fig. 1a) [13]. These neurons are poised to detect cuticle folding on the dorsal side of the animal. In addition, dorsal and ventral bipolar dendrite md neurons (dbd and vbd, respectively) extend in an anterior-posterior direction and at least dbd extends along internal connective tissue [12]. By contrast, neuron dmd1 extends an atypical thick dendrite from the body wall to the internal intersegmental nerve (ISN; [15]), which lies along the muscle layer, suggesting that this proprioceptor could be poised to detect muscle dynamics.

Imaging of dorsal class I neurons revealed that ddaE and ddaD dendrites compress in an accordion-like fashion as the peristaltic wave enters each segment, and flatten as the wave passes (Fig. 2a-a’’’; supplemental movie 4). Quantitative analysis of these dynamics shows that maximum folding of posterior ddaE dendrites occurs before maximum folding of anterior ddaD dendrites (Fig. 2b), in synchrony with the posterior to anterior progression of peristaltic waves. Like vpda, calcium dynamics revealed increases in dorsal cI activity during segment contraction (Fig. 2 c-d, supplemental movie 5, Fig. S2). Responses of posterior ddaE and anterior ddaD neurons occurred in succession during contraction, with ddaE responding just before ddaD (Fig. 3a), corresponding to the lag in dendrite folding. These data suggest that cellular calcium activity is a result of dendritic folding in all class I neurons. As a control, larvae were imaged under compression with a glass coverslip that prevented physical folding of the ddaD dendrites (Fig. S3). In this case, no increases in ddaD calcium activity were seen during forward crawling, consistent with dendritic folding driving calcium activity. This same compression did not prevent dendritic folding in ddaE neurons, and accordingly this cell type continued to show activity during crawling. This result further highlights the importance of imaging freely crawling larvae for characterization of locomotion, since physical restraint itself appears to influence proprioceptive signaling.

**Figure 2.**
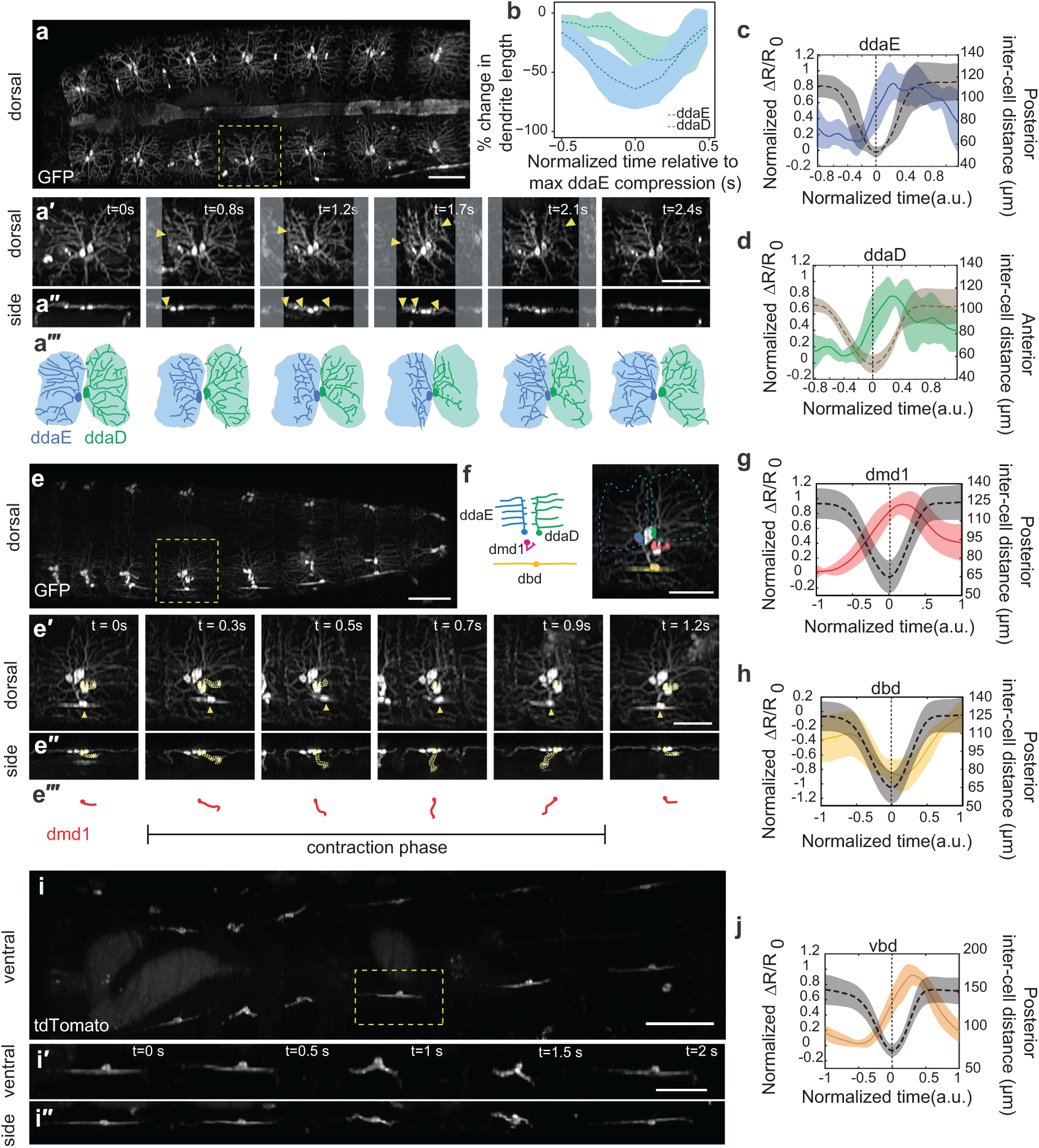
Proprioceptors with diverse morphologies showed distinct activity patterns during forwards crawling. For (**a**, **e** and **i**), images show representative SCAPE MIPs over a 80-95 μm depth range from a 160-165μm deep volume (to exclude gut autofluorescence, square root grayscale). Yellow box indicates neurons examined in time lapse sequences below, shown in both ventral and side views. **a, e** and **i** relate to supplemental movies (4-5), (6-7) and (8) respectively. **c**,**d**,**g**,**h** and **j** show mean (± s.d.) calcium response (solid line) of each proprioceptive cell type’s soma during segment contraction. Calcium response is quantified as the fractional change in the GCaMP / tdTomato fluorescence ratio ΔR/R_0_ (see methods). Mean (± s.d.) inter-cell distance is plotted with a dashed line (see methods). Maximal contraction is set at ‘t=0s’ for each recording. To compare neurons in animals crawling at different speeds, the time window and the amplitude of ΔR/R_0_ of each trace were normalized and interpolated across events (see methods). Sample sizes: **c)** 4 animals, n=5 cells, 5 events (1,1,1,2), **d)** (4 animals, n=5 cells, 5 events), (**d&h**) (n=4 animals. n=8 cells, 16 events), **j)**(*n*=*4animals*. *n*=*14 cells*, *14 events*). Fig. S2 shows GCaMP and tdTomato images as well as single-neuron GCaMP activity from all genotypes. **(a’-a’’)** shows a representative *221*-*Gal4*, *UAS*-*CD4tdGFP* larva during crawling with ddaD and ddaE neurons visible on the dorsal side. (**a’’)** Arrowheads in time-lapse indicate regions of dendrite folding. (**a’’’)** Tracing of neurons in **a**’, where shaded areas represent dendritic field territory at t=0s before folding. (**b)** Mean ± standard deviation (s.d.) percent change in dendritic length along the anteroposterior axis (a measure of dendritic folding) of ddaD (n= 10 cells) and ddaE (n= 10 cells) from *221*-*Gal4*, *UAS*-*CD4tdGFP* (2 animals) and *410*-*Gal4*, *UAS*-*CD4tdtomato* (2 animals) during forward crawling. (See Fig. S3 for changes to dendrite folding and activity in compressed conditions). (e) SCAPE imaging of *10D05*-*Gal4*, *UAS*-*CD4tdGFP* larva dorsal side during crawling, to show dorsal cluster dendrite dynamics. **(*e*’-*e*’’)** top and side views in which dmd1 (arrowhead) cell body and dendrite bundle is traced with a dashed yellow line – tracing shown in **e’’’. (f)** Schematic and inset image showing neurons imaged together in (c). Inset image shows pseudo-colored neurons. ddaE (blue), ddaD (green), dbd (yellow), dmd1 (pink). Dashed line represents outline of ddaE and ddaD dendritic fields. (**i**) SCAPE imaging of *1129*-*Gal4*, *20XUAS*-*IVS*-*GCaMP6f* (x2), *UAS*-*CD4*-*tdTomato* larva ventral side during crawling. All bounds represent s.d. Posterior is to the left for all images. Scale bar=100 μm in **a**, **e**, **i**, scale bar=50 μm in **a**’, **a**’’, **e**’, **e**’’, **f**, **i**’, **i**’’.

**Figure 3.**
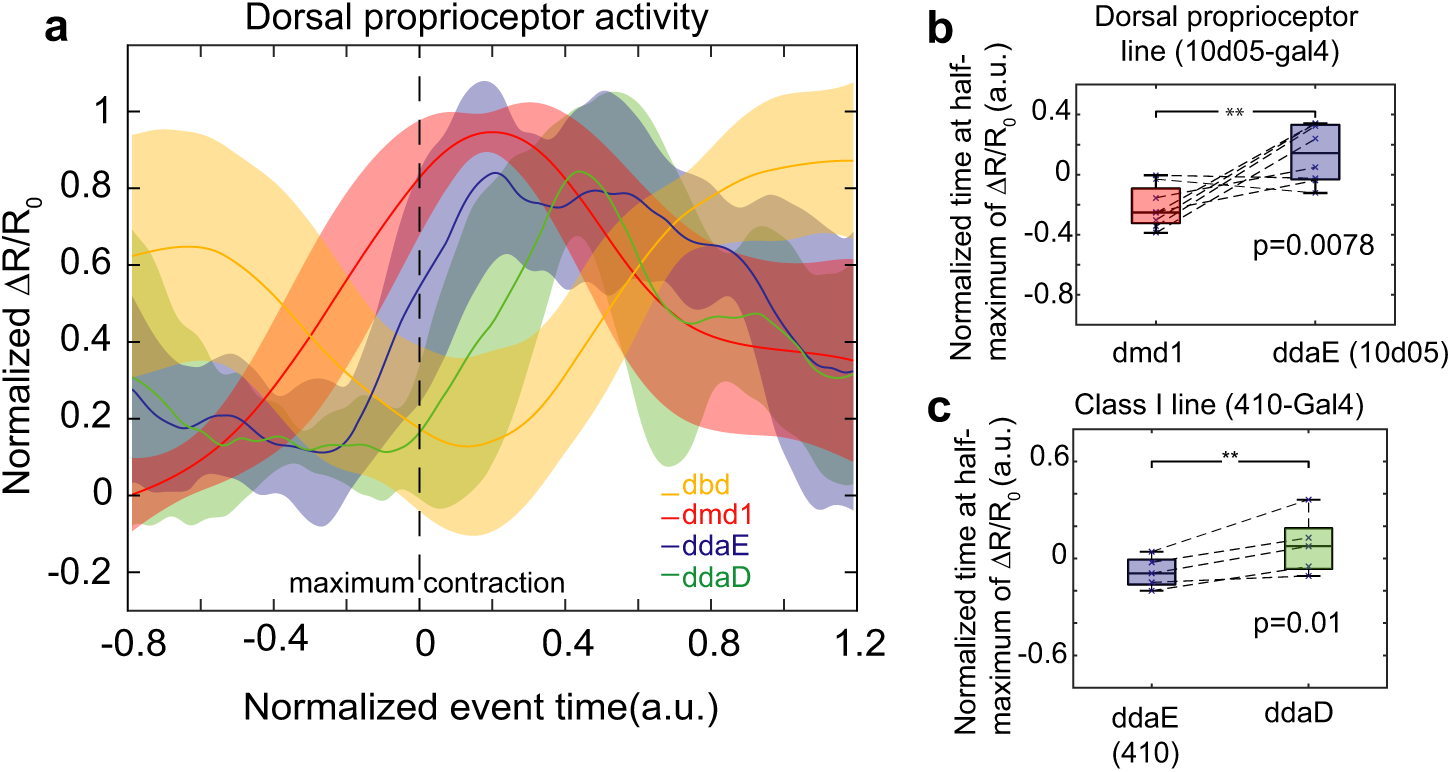
Each proprioceptor type is activated sequentially during segment contraction. (**a**) Mean calcium response of dbd, dmd1, ddaE, and ddaD during segment contraction. Data is same as shown in Fig. 2 c-d, g-h. Data for ddaE and ddaD is from *410*-*Gal4*, *20XUAS*-*IVS*-*GCaMP6f* (x2), *UAS*-*CD4*-*tdTomato* animals. Data for dbd and dmd1 is from *10D05*-*Gal4*, *20XUAS*-*IVS*-*GCaMP6f* (x2), *UAS*-*CD4*-*tdTomato* animals. Data is aligned using time at maximum contraction (as measured by distance between ddaE and homologous ddaE in posterior segment), which is set at ‘t=0s’ for each recording. The amplitudes of ΔR/R_0_ were normalized and interpolated across events. s.d is plotted as a ribbon. **(b)** To test the lag between dmd1 and ddaE activity, we compared the time at half-maximum calcium activity from *10D05*-*Gal4*, *20XUAS*-*IVS*-*GCaMP6f* (x2), *UAS*-*CD4*-*tdTomato* (n=4 animals, n=8 cells, 8 events). The central mark is the median, the edges of the box are the 25th and 75th percentiles, the whiskers extend to the most extreme data points. ddaE activity occurs significantly later than dmd1 activity (p=.0078) by single tailed paired t-test. **(c)** To test the lag between ddaE and ddaD activity, we compared the time at half-maximum calcium activity from *410*-*Gal4*, *20XUAS*-*IVS*-*GCaMP6f* (x2), *UAS*-*CD4*-*tdTomato* animals (4 animals, n=5 cells, 5 events). Note that this ddaE data set is distinct from the set in (**b**). Data is depicted as in (b). ddaD activity occurs significantly later than ddaE activity (p=0.01) by single tailed paired t-test.

The remaining three proprioceptor types have relatively internal locations, and complex 3D motion paths during crawling. We leveraged SCAPE’s high-speed volumetric imaging capabilities to capture the dynamics of these dmd1, dbd (using *10D05*-*Gal4*, Fig. 2e-f, supplemental movies 6-7, Fig. S2), and vbd (using *1129*-*Gal4*, Fig. 2i-i’’, supplemental movie 8, Fig. S2) proprioceptors during crawling. Imaging revealed 3D movements of dmd1 dendrites during muscle contraction. Prior to the contraction wave, the dendrite bundle was slack and coiled (Fig. 2e’-e’’’), then as the segment muscles contracted, the dendrite bundle stretched anteriorly and was then pulled deeper into the animal (Fig. 2e’-e’’’). As the segment relaxed, the bundle swung posteriorly, and then returned to a coiled position. GCaMP imaging revealed increases in calcium activity in dmd1 during segment contraction, as the dendrite bundle stretched (Fig. 2g).

dbd and vbd dendrites folded as the segment contracted (Fig. 2e’ arrowhead, Fig. 2i-i’’). For dbd, GCaMP fluorescence peaked during segment stretch (Fig. 2h), consistent with previous electrophysiology results in a dissected prep [21]. However, in contrast to dbd, GCaMP fluorescence in vbd peaked during segment contraction (Fig. 2j). Thus, two proprioceptors with similar morphologies and dendrite dynamics can show distinct responses during crawling.

We next tracked groups of co-labeled sensory neurons to directly compare the timing of activity in dorsal proprioceptive neurons. We found that each cell type is activated sequentially during segment contraction (Fig 3). dbd is most active in a stretched or relaxed segment. Then as the segment contracts, dmd1 increases activity first during the initial muscle contraction, followed by the cI neurons (ddaE and ddaD) as the cuticle folds during segment shortening (Fig. 3a).

Comparison of the time at half-maximum calcium activity in *410*-*Gal4* animals confirmed that ddaD activity was significantly delayed relative to ddaE during forward peristalsis (Fig. 3c; p=0.01 by single tail paired t-test). We also confirmed the lag between dmd1 and ddaE. To do this, we a used calcium activity data from *10D05*-*Gal4*, which labels all dorsal proprioceptors (Fig. 2e-f), to directly compare timing of activity of cells within the same segment. Comparison of half-maximum calcium activity confirmed that ddaE activity occurs significantly later than dmd1 activity during forward crawling (Fig. 3b, p=0.0078 by single tail paired t-test).

These data together suggest that the unique dendrite morphology of each proprioceptor tunes each cell to respond at different times during the locomotion cycle, potentially coding for different positions. This challenges the notion that some proprioceptors are redundant, and suggests that they provide a continuum of cell-type specific encoding during movement.

### Proprioceptor activity simultaneously codes for head turning and retraction

In addition to imaging simple forward crawling, our SCAPE imaging experiments captured a wide range of movements during exploration. To examine how proprioceptive activity might provide feedback during more complex body movements, we mounted the larvae in a small arena bounded by agarose. With this setup, we were able to track and extract dorsal sensory activity during exploratory head movements.

In many cases, we observed typical exploration behavior, in which larvae bend their bodies asymmetrically to turn left or right, while concurrently retracting the head backwards (Fig. 4a-a’’’; supplemental movie 9). These behaviors are not always synchronized, which leads to a somewhat complex motion path for each neuron and segment compared to locomotion. However, SCAPE recordings confirmed that even for this less-stereotyped motion, cI activity increases continue to be associated with segment contraction. In the dataset shown in Fig. 4, we focused on ddaD and ddaE within thoracic segment T3 (termed D1 and E1), and ddaD within abdominal segment A1 (termed D2), since these cells were distorted by the observed exploratory movements. Calcium activity in these cells was confirmed to correlate with the distance between ipsilateral D1 and D2 neurons (termed inter-cell distance) (Fig. 4b-c; supplemental movie 9-10, and supplemental table 1 for further discussion of E values). Signals in ddaE were found to be lower than the ddaD signals, which we reason to be due to the lesser folding of ddaE dendrites during this behavior (Fig. 4g).

**Figure 4.**
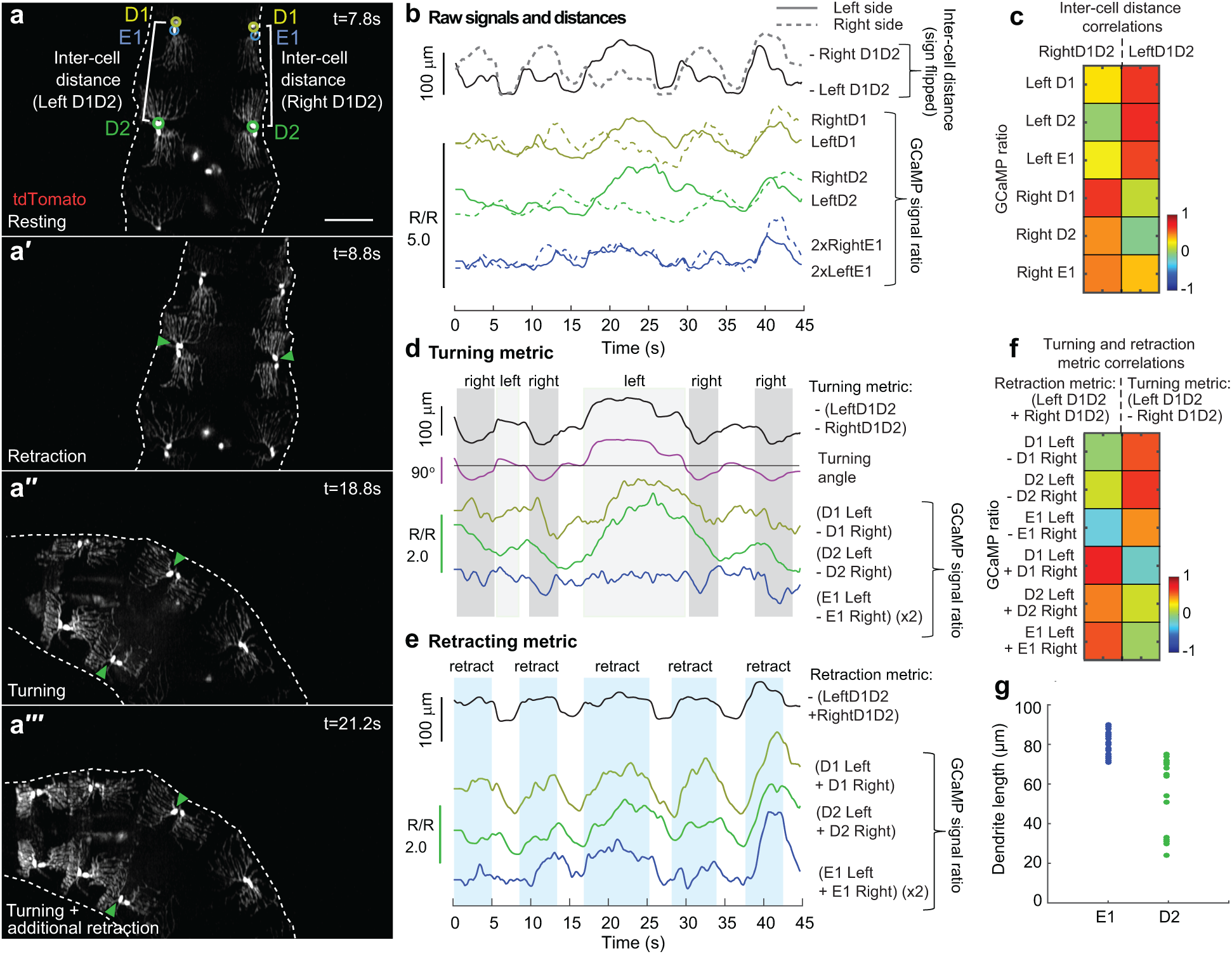
Dorsal proprioceptor activity simultaneously codes for head turning and retraction. **(a-a’’’)** SCAPE imaging of dorsal cI neurons labeled with *410*-*Gal4*, *20XUAS*-*IVS*-*GCaMP6f* (x2), *UAS*-*CD4*-*tdTomato*, dorsal side, during exploration behavior. Anterior is at the top of all images. Td-tomato channel shown. Maximum Intensity Projection (MIP) over an 80μm depth range from a 140μm deep volume. Calcium activity was quantified from circled neurons, ddaD (D1, yellow-green) and ddaE (E1, blue) in segment A1, and ddaD (D2, green) in segment A2. Inter-cell distance (white brackets) is quantified as distance between ipsilateral D1 and D2. Representative behaviors are shown including resting (**a**), retraction only (**a’)**, and turning **(a’’-a’’’)**. Turning showed different levels of retraction (e.g. **(a’’’)** shows more retraction than **(a’’)**). Supplemental movies 9-10 show this dataset. Scale bar=100 μm. All images are shown on a square root grayscale. **(b)** Calcium activity (measured as the ratio of GCaMP to tdTomato fluorescence) in left (solid) and right (dashed) D1, D2 and E1neurons are compared to inter-cell distance (black). The sign is flipped on inter-cell distance measurements, so larger values represent shorter distances. ddaE activity was smaller than ddaD activity, so E1 data is shown at 2X. **(c)** Depicts correlation coefficients between the variables shown in **b**, demonstrating positive correlations with ipsilaterally-paired (e.g. left-left) inter-cell distances and calcium signals, but near-zero correlation between contralateral measures for most cells. See supplemental table 1 for correlation values and correlations with time shifts, which show stronger correlations. **(d)** Plots of the difference between calcium activity between contralateral cells compared to the difference between contralateral inter-cell distances (our turning metric) and the calculated angle of the D1 segment. **(e)** Plots of the sum of calcium activity between contralateral cells and the sum of contralateral inter-cell distances (our retraction metric). **(f)** Depicts correlation coefficients between the variables shown in **d** and **e**. Strong correlations are seen between retraction and contralateral calcium signal sums, with no correlation to turning, while turning angle is well correlated to contralateral calcium signal differences, with no correlation to retraction. See supplemental table 2 for values and correlations with time shifts, which show stronger correlations. **(g)** The distribution of dendritic lengths (a measure of folding) during head exploration for ddaE and ddaD cells demonstrates reduced deformation of ddaE neurons, explaining the observed smaller amplitude calcium dynamics compared to ddaD neurons in **b-e**. 10 frames were sampled throughout recording, representing resting, retraction, and turning behavior.

This data during a complex movement sequence gave us the opportunity to question whether the seemingly complex calcium signals in each neuron could be interpreted in terms of their encoding of turning or head retraction independently. Based on modeling and analysis of the motion paths of the cells (see supplemental appendix), we noted that subtracting the D1-D2 intercell distance time-course for the left neuron pair from the time-course of the distance between the right neuron pair effectively cancelled out the effect of head retraction (which is symmetric on both sides of the body). The resulting differential distance (termed turning metric) was found to closely match the calculated time-varying angle of the segment, independent of retraction. Furthermore, informed by our model which assumes rigid coupling between the left and right cells, we found that adding the left D1-D2 inter-cell distance to the right D1-D2 inter-cell distance cancelled out the effects of turning (which has a reciprocal effect on left and right inter-cell distance). The result provides a measure (termed retraction metric) that represents only head retraction dynamics.

Following the same logic, we computed the difference and summation of GCaMP6f signals extracted from proprioceptor pairs on the left and right sides of the body. Remarkably, we found that the difference between calcium signals on the left versus right side correlates well with turning angle, especially for ddaD cells (Fig. 4d, f). Similarly, the sum of the calcium signals from the left and the right sides correlates strongly with our retraction metric for each cell type (D1, E1, D2) (Fig. 4e, f). Correlation values are even stronger when delays are incorporated into comparisons (Supplemental tables 1-2).

This data indicates that proprioceptor activity represents the combination of simultaneous head turning and retraction, and that calcium responses to each behavior add linearly. Furthermore, our data suggests that the larva CNS could computationally compare left versus right side activity to distinguish turning independently from head retraction.

## Discussion

This study demonstrates a new approach for live volumetric imaging of sensory activity in behaving animals, leveraging an optimized form of SCAPE microscopy [9, 10]. We used this methodology to examine the activity patterns of a heterogeneous collection of proprioceptive neurons during crawling, as well as during more complex movements such as head turning and retraction, to determine how larvae sense body shape dynamics. Imaging revealed 3D distortion of proprioceptive dendrites during crawling, and GCaMP activity that occurred coincident with dendritic deformations. Most neurons (all cI neurons, dmd1, and vbd) increased activity during segment contraction. In contrast, dbd neurons increased activity during segment stretch or relaxation, which is consistent with previous electrophysiological recordings of dbd in a dissected preparation [21].

The temporal precision of SCAPE revealed that during forward crawling, different proprioceptors exhibit sequential onset of activity, associated with their unique dendrite morphologies and movement dynamics, suggesting that proprioceptors monitor different features of segment deformation. The complementary sensing of compression versus stretch in cI/dmd1/vbd versus dbd neurons provides an additional measure of movement that is conceptually similar to the responses of Golgi tendon organs versus muscle spindles in mammals [22]. Combined, these results indicate that this set of proprioceptors function together to provide a continuum of sensory feedback describing the diverse 3D dynamics of the larval body.

Prior work suggested that the proprioceptors studied here have redundant functions during forward crawling because silencing different subsets caused similar behavioral deficits, namely slower crawling [14]. Slow locomotion may be a common outcome in a larva that is lacking in part of its sensory feedback circuit, yet our results suggest that each cell type has a unique role. Our demonstration of the varying activity dynamics of proprioceptors during crawling and more complex movements suggests that the larva utilizes feedback from a combination of these unique sensors to infer diverse aspects of speed, angle, restraint and overall body deformation. This feedback system is likely to be important for a wide range of complex behaviors, such as body bending and nociceptive escape.

How can an understanding of proprioceptor activity patterns inform models of sensory feedback during locomotion? Electron microscopic reconstruction has shown that ddaD, vbd, and dmd1 proprioceptors synapse onto inhibitory PMSI pre-motor neurons (A02b) [23], which promote segment relaxation and anterior wave propagation [24]. Thus, activity of these sensory neurons may signal successful segment contraction and promote forward locomotion, in part by promoting segment relaxation. Furthermore, vpda neurons provide input onto excitatory premotor neurons A27h, which acts through GDL interneurons to inhibit contraction in neighboring anterior segments, thereby preventing premature wave propagation [25]. In this way, vpda feedback could contribute to proper timing of contraction in anterior segments during forwards crawling. In contrast to other proprioceptors, dbd neurons are active during segment stretch. Their connectivity also tends to segregate from contraction sensing neurons [23, 26] and understanding how the timing of this input promotes wave propagation is an important future question. Our dynamic recordings of the function of these neurons during not just crawling, but also exploration behavior provides essential new boundary data for testing putative network models derived from this anatomical roadmap.

SCAPE’s high-speed 3D imaging capabilities enabled 10 VPS imaging of larvae during rapid locomotion. Fast volumetric imaging not only prevented motion artifacts, but it also revealed both the 3D motion dynamics and cellular activity associated with crawling behavior. SCAPE’s large, 1 mm wide field of view allowed multiple cells along the larva to be monitored at once, while providing sufficient resolution to identify individual dendrite branches. Because SCAPE data is truly 3D, dynamics could be examined in any section or view. Additionally, fast two-color imaging enabled simultaneous 3D tracking of cells, monitoring of GCaMP activity, and correction for motion-related intensity effects. Our demonstration that larvae that are compressed during crawling exhibit altered dendrite deformation, and thus altered proprioceptive signaling (Fig S2) underscores the benefit of being able to image unconstrained larvae, volumetrically in real-time. Furthermore, rapid volumetric imaging allowed for the analysis of sensory responses during non-stereotyped, exploratory head movements in 3-dimensions, revealing a combinatorial code for proprioceptor encoding of complex, simultaneous movements. This finding also demonstrates the quantitative nature of SCAPE data and its high signal to noise, which enabled real-time imaging of neural responses without averaging from multiple neurons.

Here, we provide an example of how high-resolution, high-speed volumetric imaging enabled investigation of the previously intractable question of how different types of proprioceptive neurons encode forward locomotion and exploration behavior during naturalistic movement. Imaging could readily be extended to explore a wider range of locomotor behaviors such as escape behavior, in addition to other sensory modalities such as gustation and olfaction. Detectable signals reveal rich details including the firing dynamics of dendrites and axonal projections during crawling. Waves of activity in central neurons within the ventral nerve cord can also be observed. We expect that the *in vivo* SCAPE microscopy platform utilized here could ultimately allow complete activity mapping of sensory activity during naturalistic behaviors throughout the larval CNS. Using SCAPE, it is conceivable to assess how activity from proprioceptive neurons modulate central circuits that execute motor outputs, which will provide critical information for a dissection of the neural control of behavior with whole animal resolution.

## Acknowledgments

We thank Dr. Gary Struhl for discussions and Dr. Oliver Hobert for reading an earlier version of the manuscript. We thank Dr. Grace Ji-eun Shin for suggestions on the manuscript. We thank the Clandinin lab for sharing the InSITE collection, Katarzyna Rojek and Dr. Dietmar Schmucker for sharing expression data for *410*-*Gal4*, and Dr. Matthew Bouchard for SCAPE development and early assistance with experiments and Thomas Bernhardt for initial work on the cell tracking code for SCAPE analysis. Ying Ma for assistance with statistical analysis. We thank Dr. Dan Tracey for communication of results prior to publication. This work was supported by NIH 5R01NS061908 (W.B.G.), NIH BRAIN 5U01NS094296 and UF1NS108213 (E.M.C.H., W.B.G.), the McKnight Foundation (W.B.G.), DoD, MURI W911NF-12-1-0594 and the Simons Foundation Collaboration on Global Brain (E.M.C.H). R.V. was supported by NIH 5T32HD007430-18 and an NSF predoctoral fellowship.

## Author contributions

Conceptualization, W.B.G., E.M.C.H.; Methodology, R.V., W.L., V.V., E.M.C.H., W.B.G.; SCAPE development and optimization: V.V., W.L., E.M.C.H; Software, W.L., E.M.C.H.; Validation, R.V. W.L., E.M.C.H., W.B.G.; Formal analysis, R.V., W.L., E.M.C.H., W.B.G.; Investigation, R.V., W.L., A.S., V.V., E.M.C.H., W.B.G.; Writing – Original Draft, R.V., W.L., E.M.C.H., W.B.G.; Writing – Reviewing & Editing, R.V., W.L., A.S., E.M.C.H., W.B.G.; Visualization, R.V., W.L., E.M.C.H., W.B.G.; Supervision, E.M.C.H., W.B.G. Project Administration, E.M.C.H., W.B.G., Funding Acquisition, E.M.C.H., W.B.G.

## Methods

### Contact for Reagent and Resource Sharing

Further information and requests for resources should be directed to and will be fulfilled by the Lead Contact, Elizabeth Hillman. Request for *Drosophila* reagents should be directed to Wesley Grueber, Corresponding Author.

### Experimental Model and Subject Details

To image cI da neurons we used *221*-*Gal4*, *UAS CD4*-*tdGFP* and *IT*.*410*-*Gal4*, *20XUAS*-*IVS*-*GCaMP6f* (2 copies), *UAS*-*CD4*-*tdTomato*. To image GFP dynamics as a control, we used *IT*.*410*-*Gal4*, *20XUAS mCD8::GFP*, *UAS*-*CD4*-*tdTomato*. To image all dorsal proprioceptors, we used *10D05*-*Gal4*, *20XUAS mCD8::GFP* or *10D05*-*Gal4*, *20XUAS*-*IVS*-*GCaMP6f (x2) (2 copies)*, *UAS*-*CD4*-*tdTomato*. To image vbd we used *IT*.*1129*-*Gal4*, *20XUAS*-*IVS*-*GCaMP6f (x2) (2 copies)*, *UAS*-*CD4*-*tdTomato*. 2^nd^ instar larvae were used for all imaging experiments, except 3^rd^ instar larvae were used for imaging of compressed animals.

### Method Details

#### SCAPE Image acquisition

High-speed volumetric imaging of crawling larvae was performed using a custom swept confocally aligned planar excitation (SCAPE) microscope extended from designs described in [9, 10]. Briefly, high speed 3D imaging is achieved by illuminating the sample with an oblique light sheet through a high NA objective lens [27]. Fluorescence light excited by this sheet (extending in y-z’) is collected by the same objective lens (in this case an Olympus XLUMPLFLN 20XW 1.0 NA water immersion objective with a 2mm working distance). A galvanometer mirror in the system is positioned to both cause the oblique light sheet to scan from side to side across the sample (in the x direction, without a change in the angle of the sheet) but also to descan returning fluorescence light. This optical path results in an intermediate, descanned oblique image plane which is stationary yet always co-aligned with the plane in the sample that is being illuminated by the scanning light sheet. Image rotation optics and a fast sCMOS camera (Andor Zyla 4.2) are then focused to capture these y-z’ images at over 1000 frames per second as the sheet is scanned in the sample in the x direction. Data is then reshaped into a 3D volume by stacking successive y-z’ planes according to the scanning mirror’s x-position. All other system parts including the objective and sample stage are stationary during high speed 3D image acquisition, including the primary objective lens and sample stage. No image reconstruction procedures besides correction for the sheet’s oblique angle were used for data shown in this study.

SCAPE’s primary objective was configured in an inverted arrangement for the ventral side imaging and an upright arrangement for the dorsal side imaging. Dual-color imaging was achieved using a custom-built dual color image splitter in front of the sCMOS camera. 488 nm excitation (<5 mW at the sample, Coherent OBIS) was used to excite fluorescence in both channels, with 525/45 nm and 600/50 nm emission filters in the green and red emission channels respectively. The system’s camera frame rate to read 150-200 rows (corresponding to oblique depths along z’) was 1000-1300Hz, with an x-scanning step size of 2~3μm to achieve 10 volumes per second imaging over a field of view of 1000 × 250 × 195 μm (y-x-z)(scan parameters varied for different trials, based on the size difference of each larva).

To image forward crawling, 2nd instar larvae were chosen to image multiple segments at once. Larvae were imaged while positioned within a 300μm wide water-filled channel bounded by FEP spacers and covered by a 40mm × 24mm cover glass. When imaging the ventral side, the channel was positioned on a 50mm × 24mm cover glass, when imaging the dorsal side, the channel was positioned on a glass slide. Each trial acquired data for up to 120 seconds or was terminated earlier if the larva crawled to the end of the channel. A manual translation stage aligned along the FEP channel axis was used to keep the larva in the field of view during the acquisition as needed.

To image compressed animals, we reproduced typical conditions for confocal microscopy imaging of larvae. As such, 3rd instar larvae were positioned in a 50:50 mixture of halocarbon oils 27 and 700 to enhance compression with an overlying coverslip. We ensured that cross sections of the animal showed body compression during imaging (Fig. S3).

To image complex head movements (turns and retractions) during exploration behavior, we constrained 2^nd^ instar larvae in a small water-filled arena bounded by 10% agarose, with a coverslip on top. The size of the arena is about 1000μm by 500μm and it was made on a ~200μm thick agarose pad.

All the functional calcium signal analysis used signals extracted from raw, linear-scale imaging data. However, for the visualizations of SCAPE images shown in the figures and movies, raw camera 16bit data was square root scaled to enhance the visible dynamic range to avoid display saturation and to make all components (soma and dendrites) more visible. Resulting pixel values are then shown on a linear gray, red or green colorscale without further adjustment. SCAPE data was interpolated to uniform voxels with spline smoothing for all the figures and movies, and sharpened using the imageJ function ‘unsharp mask’ (radius:1.5, weight:0.4) to enhance dendrite visualization except for Figure 1d-d’ and supplementary movies 2, 3, 5, 7, 8. None of the raw SCAPE imaging data was saturated during acquisition. Images and movies were generated using Matlab and ImageJ.

#### Cell tracking and ratiometric correction of calcium activity

For SCAPE imaging quantification, vpda, vbd and dbd neurons were first manually selected and then automatically tracked in 3D space for the duration of the run that the neuron was within the field of view, using Matlab. For other neuron types (ddaD, ddaE, dmd1), since the distance between neighboring neurons is small, automatic tracking was performed under supervision and manually corrected when needed. This 3D tracking provided behavioral information related to the animal’s physical movements, as well as fiducials for extraction of GCaMP fluorescence from the cells during movement.

For fluorescence extraction, tracking regions of interest (ROIs) were defined as the smallest rectangular 3D cube around the tracked cell that encompassed the entire cell body. Average fluorescence intensity values of GCaMP6f and tdTomato were then extracted from these ROIs for each time point. A ratio between GCaMP6f and tdTomato was calculated after subtraction of background signal to account for the motion induced intensity change for each frame (yielding the green-to-red ratio R). The average of the lowest 10% ratio values was used as the baseline (R_0_) for each ROI. The GCaMP signal reported as neural activity at each time point then corresponds to the change in this ratio from baseline (ΔR/R_0_). To demonstrate this process, raw red, green and ratiometric signals are shown in Figure S1. In addition, control measurements are shown that applied the same analysis to larvae co-expressing tdTomato and static GFP. In the GFP case, dynamic changes in ΔR/R_0_ were insignificant, confirming the sensitivity of our (ΔR/R_0_) measure to the intracellular calcium-dependent fluorescence of GCaMP.

#### Calculating segment and dendrite length dynamics

To relate calcium dynamics to segmental contraction and extension phases, changes in inter-cell distances were calculated from the tracked cell coordinates - defined as the distance between the measured neuron and a homologous neuron in the posterior or anterior segment over time. For vpda and vbd, we plotted posterior-intercell distance between the measured neuron and a homologous neuron in the posterior segment. For dbd, dmd1, ddaE, and ddaD we plotted the posterior inter-cell distance between the ddaE neuron in the same segment as the cell of interest and the homologous neuron in the next posterior segment. This allowed us to directly compare the timing of dorsal neuron activity. For some plots of ddaD activity (Fig. 2d, Fig. S3), we plotted anterior inter-cell distance between the measured neuron and the homologous ddaD neuron in the anterior segment, since this was a better proxy for dendrite folding.

To directly measure dendrite dynamics in ddaD and ddaE neurons (Fig. 2b), 2-3 cells of each type from segments A1-A6 were analyzed from 4 different animals. Dendrite length was measured as a 180 degree line from the cell body to the furthest visible dendrite. Each cell was analyzed during a different peristaltic wave.

#### Normalizing and averaging dynamics across larvae

Real-time traces of inter-cell distance and ΔR/R_0_ shown in Figures 1, S1, and S2, demonstrate the high signal to noise and repeatability of our observations. However, these traces also demonstrate that the speed of crawling can vary quite significantly between animals, in addition to the relative amplitude of (ΔR/R_0_). To provide the aggregate, average properties of neural activity against segment contraction, it was thus necessary to normalize these differences between animals. To calculate mean calcium responses (ΔR/R_0_) in relation to segment contraction, we normalized amplitude of responses across events to 1, so as not to bias the average values to cells that responded more strongly. To plot mean calcium activity (ΔR/R_0_), inter-cell distance, or dendrite measurements across animals with different crawling speeds, normalization was applied in time based on the full width at half maximum (FWHM) of the mean inter-cell distance of every contraction for each animal. 1 A.U. ranges from 0.7-2.5 seconds. To test the activity timing lag between neuron types (Fig. 3b-c), we compared the normalized time at half-maximum calcium activity.

To characterize the resting and contraction phase activity in Figure S1c, we defined the contraction phase as 2×FWHM window centered at the maximum contraction point and resting phase as the 0.5×FWHM window prior to the contraction phase. The resting phase GFP-tdTomato or GCaMP6f-tdtomato ratio value was calculated by taking the mean ΔR/R_0_ along the resting phase window. And the contraction phase GFP-tdTomato or GCaMP6f-tdtomato ratio value was calculated by taking the maximum ΔR/R_0_ over the contraction phase window.

### Quantification and Statistical Analysis

Statistical tests were performed using Matlab. We evaluated the lag between dmd1 vs. ddaE and ddaE vs. ddaD activity by single tailed paired t-test, alpha level 0.05. We evaluated the difference between the resting and contraction phases for the GFP-tdTomato and GCaMP6f-tdtomato ratios with two-tailed paired t-test. Statistical parameters reported in figure legends. All p values are represented as: ^∗^ < 0.05, ^∗∗^ < 0.01, and ^∗∗∗^ < 0.001.

### Data and Software Availability

Cell tracking, signal extraction and ratiometric correction was performed using custom Matlab scripts that will be made available upon request. The data that support the findings of this study are available from the corresponding authors upon reasonable request.

## Supplemental Information

**Figure S1.**
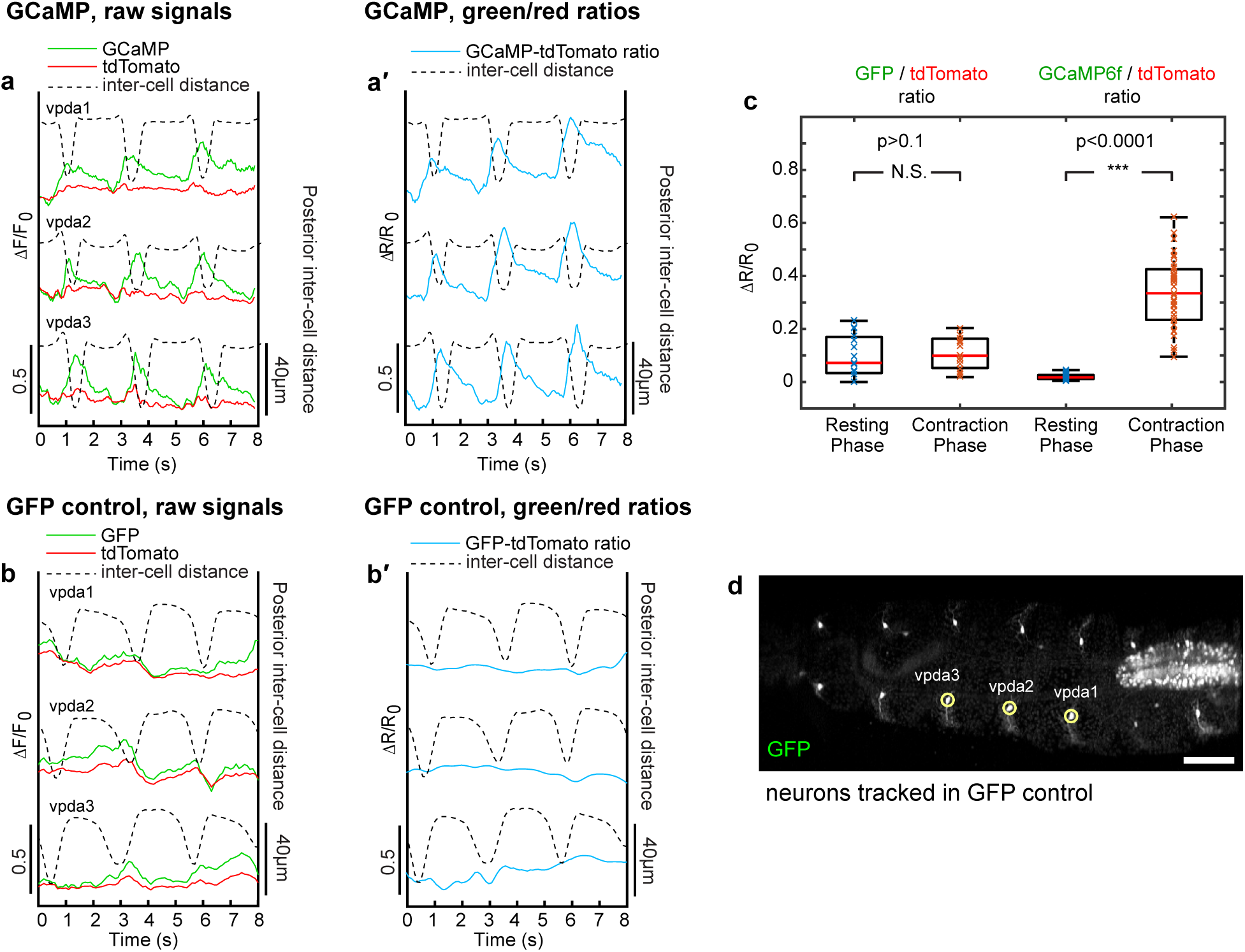
Ratiometrically measured calcium dynamics properly control for motion artifacts, related to Figure 1. **(a)** Change in fluorescence from baseline (ΔF/F_0_) in GCaMP6f (green) and tdTomato (red) during crawling in vpda neurons. Segment contraction is depicted with inter-cell distance (dashed lines). **(a’)** Change in ratio of GCaMP6f to tdTomato fluorescence (ΔR/R_0_, blue). Increases can be seen during segment contraction. **(b)** Change in fluorescence from baseline (ΔF/F_0_) in GFP (green) and tdTomato (red) during crawling in vpda neurons. Segment contraction is depicted with inter-cell distance (dashed lines). (**b**’) Change in ratio of GFP to tdTomato fluorescence (ΔR/R_0_, blue). No increase is associated with segment contraction. **(c)** Comparison of GFP-tdTomato and GCaMP6f-tdtomato ratios between resting and contraction phases (see methods). For GFP-tdTomato analysis, n= 2 animals, 7 cells, 14 events, for GCaMP6f-tdtomato analysis, n=3 animals, 22 cells, 26 events. Note that there is no difference between GFP-tdTomato ratios in the resting versus contraction phases, while there is a significant increase in GCaMP6f-tdTomato ratios during contraction (p<0.001, as measured by two-tailed t-test). (**d**) SCAPE imaging of *410*-*Gal4*, *20XUAS mCD8::GFP*, *UAS*-*CD4*-*tdTomato* animals during forward crawling, Ventral side. Imaging shows vpda neurons. GFP channel is shown. Posterior is to the left. Images are shown on a square root grayscale to reduce dynamic range for visualization of both cell bodies and dendrites. Scale bar=100μm.

**Figure S2.**
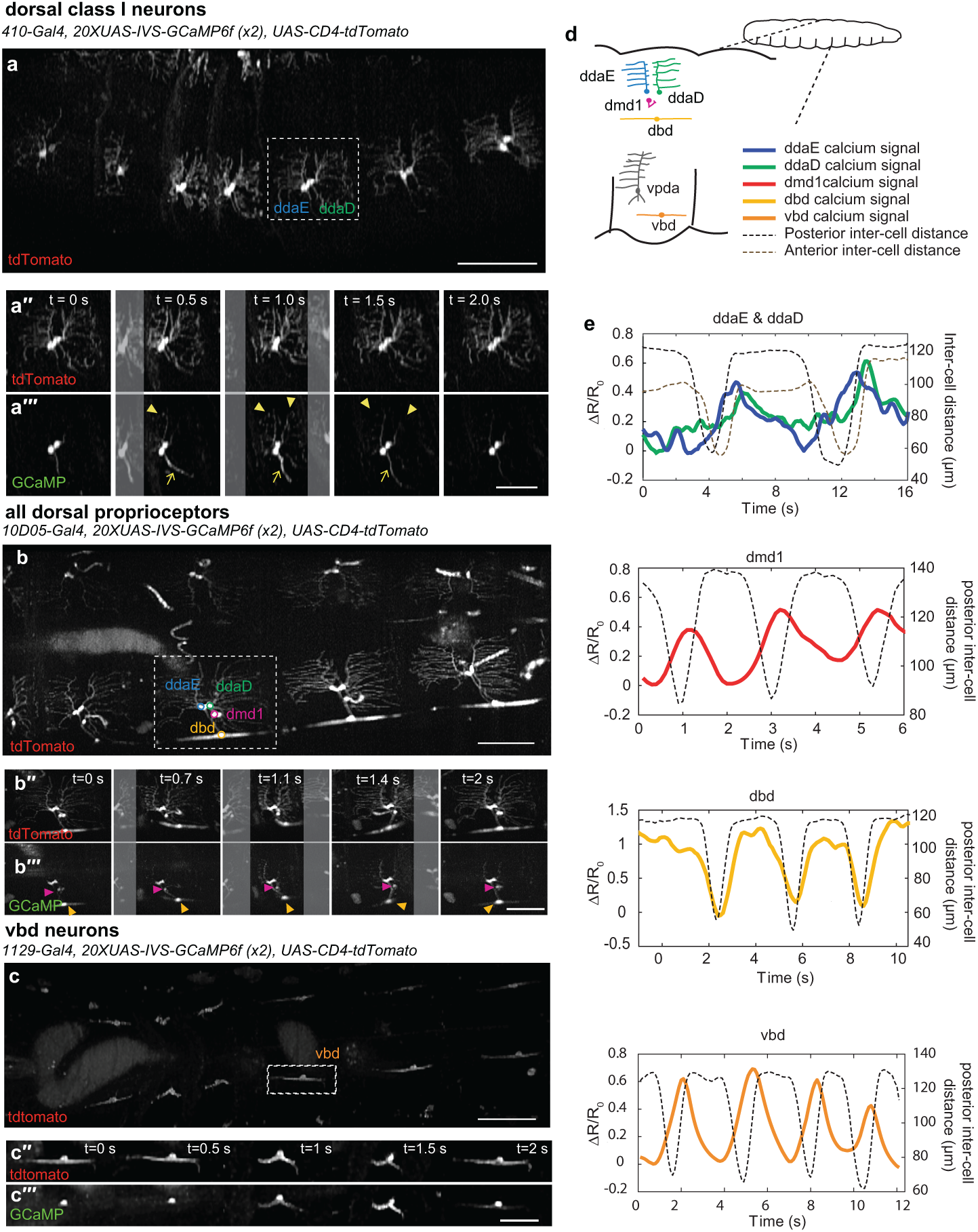
Examples of SCAPE imaging of GCaMP dynamics, related to Figure 2. Posterior is to the left for all images. For (**a-c**), images show representative SCAPE MIPs over a 35-90 μm depth range from a 160-200 deep volume (to exclude gut autofluorescence, square root grayscale). Dashed box indicates neurons examined in time lapse sequences below, shown for both tdTomato and GCaMP channels. **(a-a’’’)** SCAPE imaging of *410*-*Gal4*, *20XUAS*-*IVS*-*GCaMP6f* (x2), *UAS*-*CD4*-*tdTomato* larva. See supplemental movie 5. Arrowheads indicate increases in dendritic GCaMP6f, arrows indicate increases in axon bundle (containing both ddaD and ddaE axons). Note ddaE dendrites are active before ddaD. **(b-b’’’)** SCAPE imaging of *10D05*-*Gal4*, *20XUAS*-*IVS*-*GCaMP6f* (x2), *UAS*-*CD4*-*tdTomato* larva. See supplemental movie 7. Orange arrowhead marks dbd cell body, pink arrowhead marks dmd1 cell body. **(cc’’’)** SCAPE imaging of *1129*-*Gal4*, *20XUAS*-*IVS*-*GCaMP6f* (x2), *UAS*-*CD4*-*tdTomato* larva. See supplemental movie 8. **(d)** Schematic of larval proprioceptive system **(e)** Examples of single cell calcium activity dynamics during forward crawling. The calcium response is plotted in solid lines (quantified as ΔR/R_0_). The distance between the measured neuron and the posterior neuron (posterior inter-cell distance) is plotted in black dashed lines. The distance between the measured neuron and the anterior neuron (anterior inter-cell distance) is also plotted in brown dashed lines on the ddaD plot, since this is a better proxy for dendrite folding.

**Figure S3.**
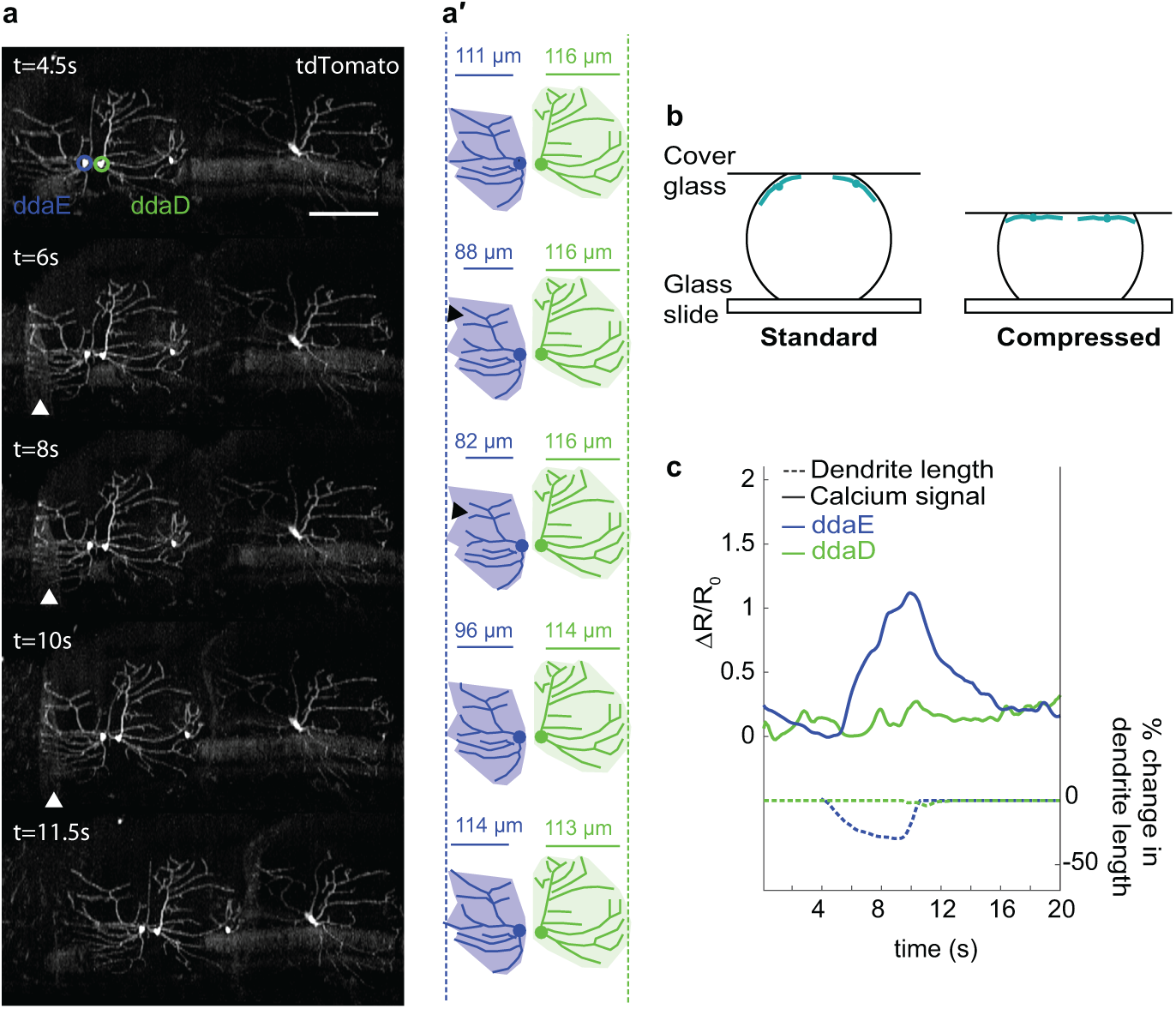
Sensory activity does not occur in the absence of dendritic folding, related to Figure 2. **(a)** Time lapse of SCAPE imaging of dorsal class I neurons labeled with *410*-*Gal4*, *20XUAS*-*IVS*-*GCaMP6f* (x2), *UAS*-*CD4*-*tdTomato*, in a compressed preparation, which prevents dendritic folding in ddaD (see **(b)** and methods). TdTomato channel is shown to depict dendrite dynamics. Larva is 3^rd^ instar. Posterior is to the left. (MIP) over a 50μm depth range from a 160μm deep volume. **(a’)** Tracing of time lapse data shown in **(a)**, posterior cells. ddaE is blue and ddaD is green. Dotted lines and shaded areas represent extent of arbor in a relaxed segment. Measurements represent dendrite length (μm), a measure of dendrite folding. Arrows denote frames with dendrite folding. Note that ddaE dendrites fold, but not ddaD. **(b)** Schematic of compressed preparation. **(c)** Calcium responses (ΔR/R_0_, solid lines) and % change in dendrite length (dotted lines) in a compressed preparation of ddaE (blue) and ddaD (green) during segment contraction. Activity correlates with dendrite folding. Scale bar=100μm.

## Supplemental Appendix Motion model for exploring larva.

We made the following assumptions based on the properties of motion of the tracked neurons D1L (left anterior ddaD), D1R (right anterior ddaD), D2L (left posterior ddaD) and D2R (right posterior ddaD) that:

1. The lava’s right and left neurons are positioned either side of a rigid bar of length ~2p
2. Turning motion is given by a time-varying rotation θ (t) of the anterior bar about the midpoint of the posterior bar.
3. The retraction motion of the larva was approximated as a time-varying distance S(t) between the midpoints of each bar.

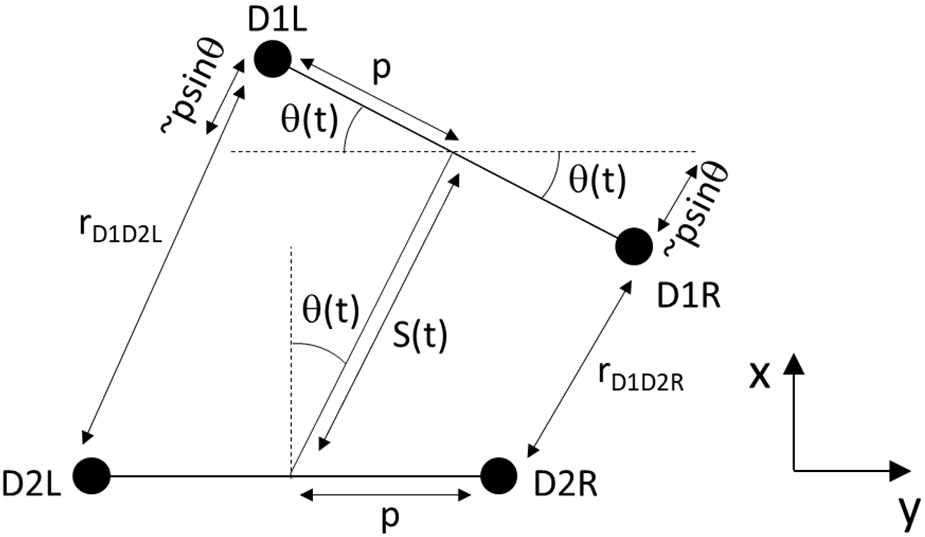

From this construction, we can see that good approximations to the distance between D1R and D2R (rD1D2R) and D1L and D2L (rD1D2L) are simply:

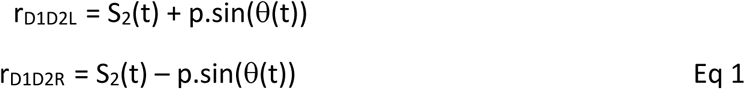

From this relationship, it is simple to see that the subtraction or summation of these two inter-cell differences are going to yield:

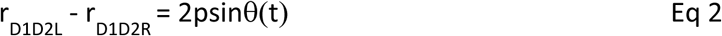

which is a pure function of turning angle, and

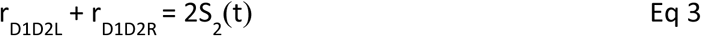

which is a pure function of time-varying head retraction.

This ‘common mode rejection’ property holds true for the positional data of the larva’s neurons, as well as the subtraction and summation of the left and right GCaMP signal extracted from neurons innervating the inter-cell space.

A more rigorous derivation notes that the precise value of the inter-cell distance is given by:

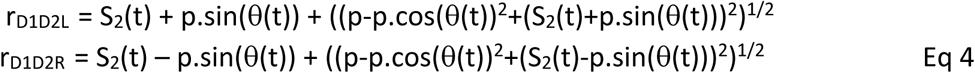

however, the difference between this and our approximation is <0.12% for the motion parameters of the larva observed.

**Supplementary Table 1:** Correlation coefficient values shown in Figure 4c, time-aligned and with a time-shift:

|  | Inter-cell distances |  |  |  |  |  |
| --- | --- | --- | --- | --- | --- | --- |
| $\Delta R/R$ signal changes: | Left D1 | Left D2 | Left E1 | Right D1 | Right D2 | Right E1 |
| Left D1D2 | 0.64 | 0.67 | 0.60 | 0.10 | -0.08 | 0.35 |
| Right D1D2 | 0.31 | -0.02 | 0.24 | 0.66 | 0.45 | 0.48 |
| With time-shift (1.2 sec for D1 and 0.45 sec for E): |  |  |  |  |  |  |
| Left D1D2 | 0.85 | 0.86 | 0.67 | 0.21 | 0.09 | 0.43 |
| Right D1D2 | 0.26 | -0.03 | 0.33 | 0.86 | 0.79 | 0.55 |

**Supplementary Table 2:** Correlation coefficient values shown in Figure 4f, time-aligned and with a time-shift:

|  | Inter-cell distances |  |  |  |  |  |
| --- | --- | --- | --- | --- | --- | --- |
| $\Delta R/R$ signal changes: | D1 Left-Right | D2 Left-Right | E1 Left-Right | D1 Left + Right | D2 Left + Right | E1 Left + Right |
| (Turning metric)<br>Left D1D2 - Right D1D2 | 0.59 | 0.64 | 0.45 | -0.20 | 0.13 | 0.02 |
| (Retraction metric)<br>Left D1D2 + Right D1D2 | -0.01 | 0.14 | -0.28 | 0.69 | 0.50 | 0.57 |
| With time-shift (1.2 sec for D1 and 0.45sec for E): |  |  |  |  |  |  |
| (Turning metric)<br>Left D1D2 - Right D1D2 | 0.82 | 0.84 | 0.44 | -0.15 | 0.16 | 0.01 |
| (Retraction metric)<br>Left D1D2 + Right D1D2 | -0.12 | 0.02 | -0.31 | 0.88 | 0.81 | 0.68 |

Shifting the neuronal calcium responses in time accounts for the phase lag in signaling compared to maximal contraction, consistent with results shown in Figure 3. Lower correlation coefficient values relating to the Right E1 neuron may relate to the more minimal compression of E dendrites (Fig 4g) and the generally smaller amplitude of right-hand turns. The E neuron pair generally appears to be more sensitive to retraction than turning, which explains the relatively similar correlation levels between Right E1 and left and right D1-D2 distances.

## Supplemental Movie Legends

**Supplemental movie 1.**
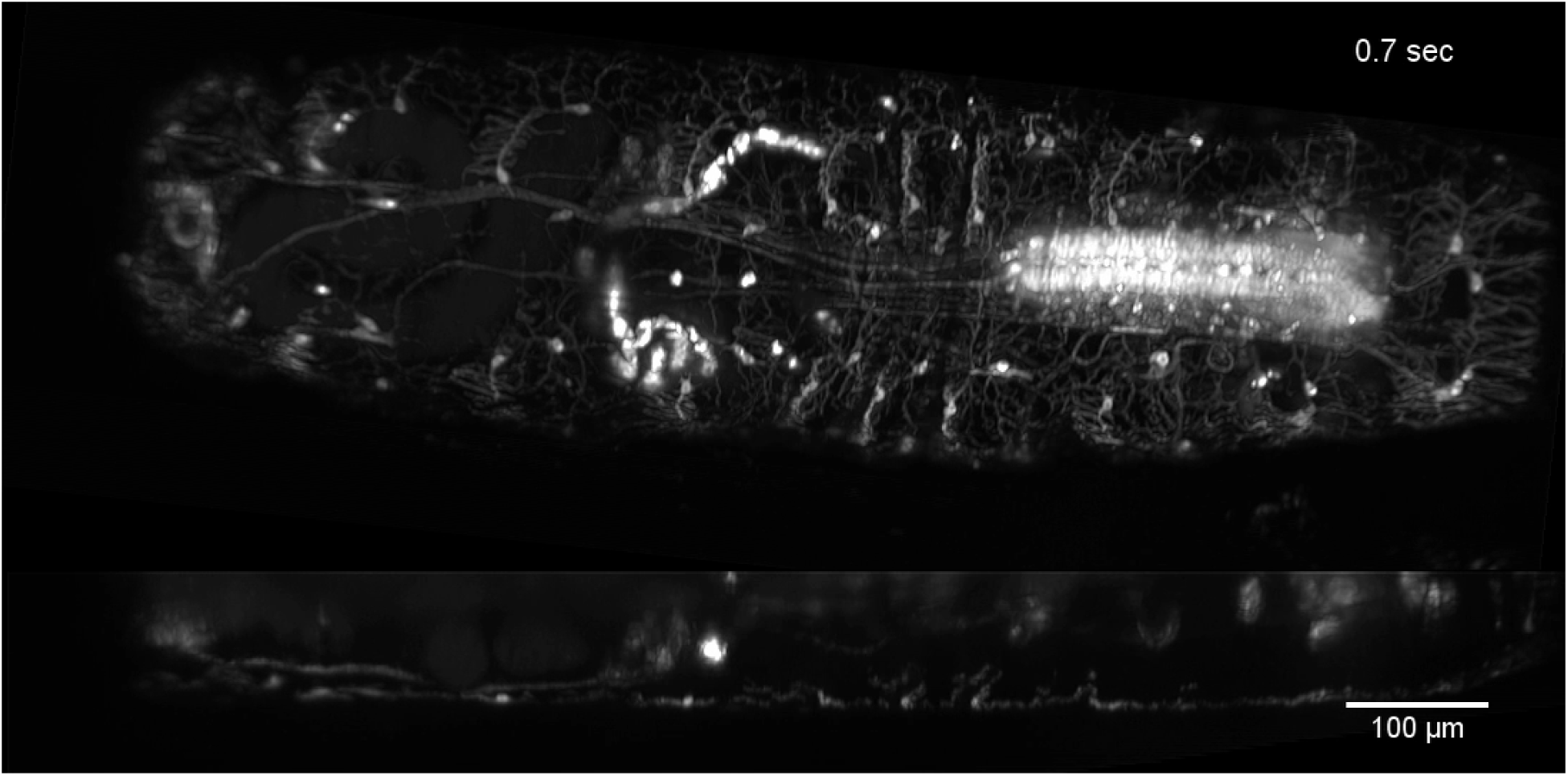
Ventral class I dendrite dynamics during crawling, GFP only. Class I dendrites (and class IV weakly) are labeled by *221*-*Gal4*, *UAS-mCD8::GFP*. Top is ventral view MIP over a 95 μm depth range from a 160μm deep volume, bottom is orthogonal view of ventral cI neuron vpda. Note that dendrites of vpda fold with each peristaltic wave. SCAPE images are shown on a square root grayscale to reduce dynamic range for visualization of both cell bodies and dendrites. Posterior is to the left. Scale bar=100μm.

**Supplemental movie 2.**
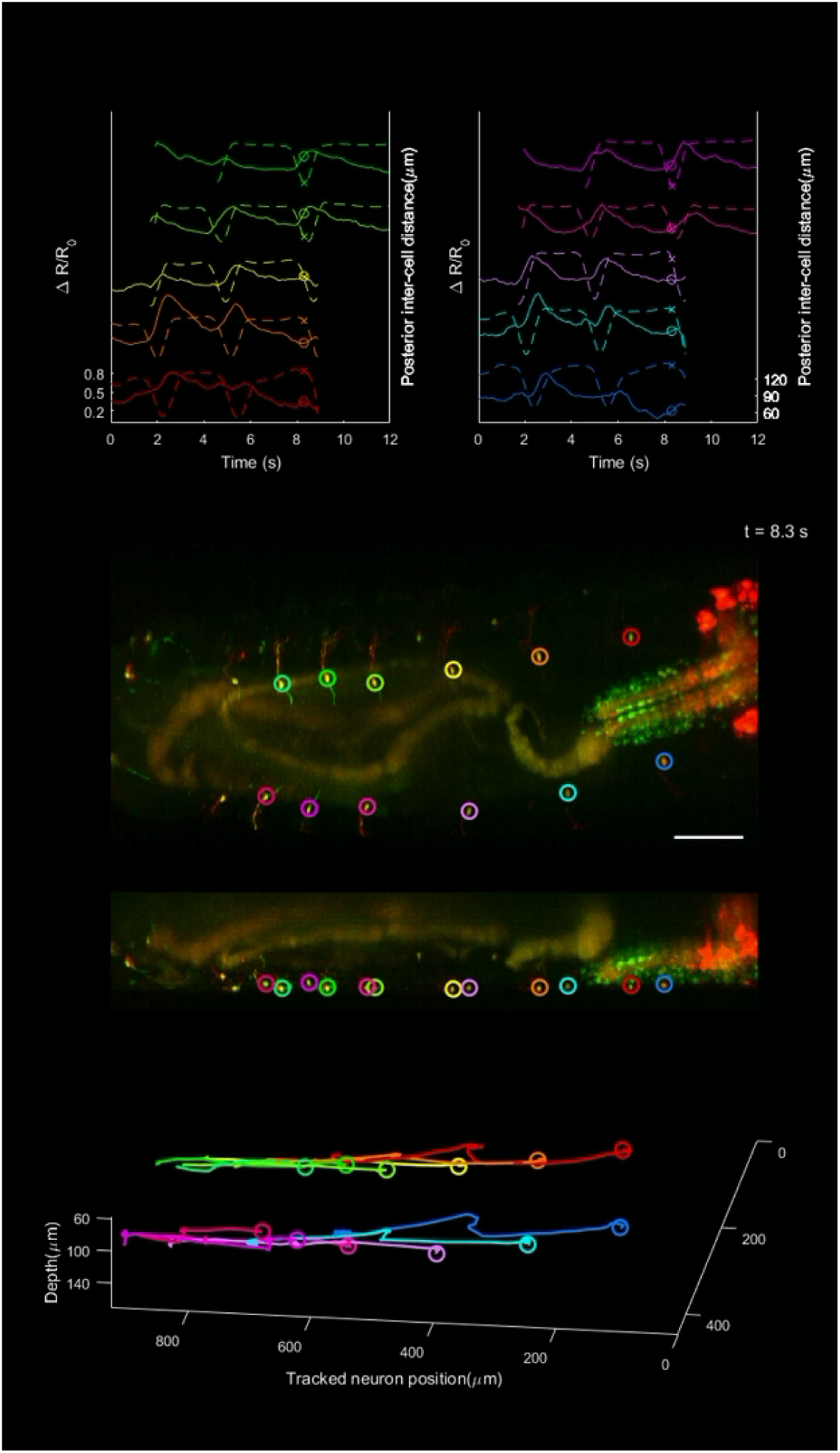
Tracking of neurons and quantification of GCaMP6f fluorescence during crawling using SCAPE microscopy. cI neurons are labeled with *410*-*Gal4*, *20XUAS*-*IVS*-*GCaMP6f* (x2), *UAS*-*CD4*-*tdTomato*. Top panels show five tracked vpda neurons each on left and right of larva. Solid line indicates GCaMP fluorescence (measured as ΔR/R_0_) and dashed lines indicate distance between tracked cell and posterior cell (inter-cell distance). Middle panels depict dual channel SCAPE imaging of a crawling larva (ventral MIP from full 168 μm deep imaging volume and an orthogonal MIP view, square root colorscale). CNS is observed at right margin of the movie. Bottom panel is the positions of 12 tracked neurons in 3D space. Posterior is to the left. Scale bar=100μm.

**Supplemental movie 3.**
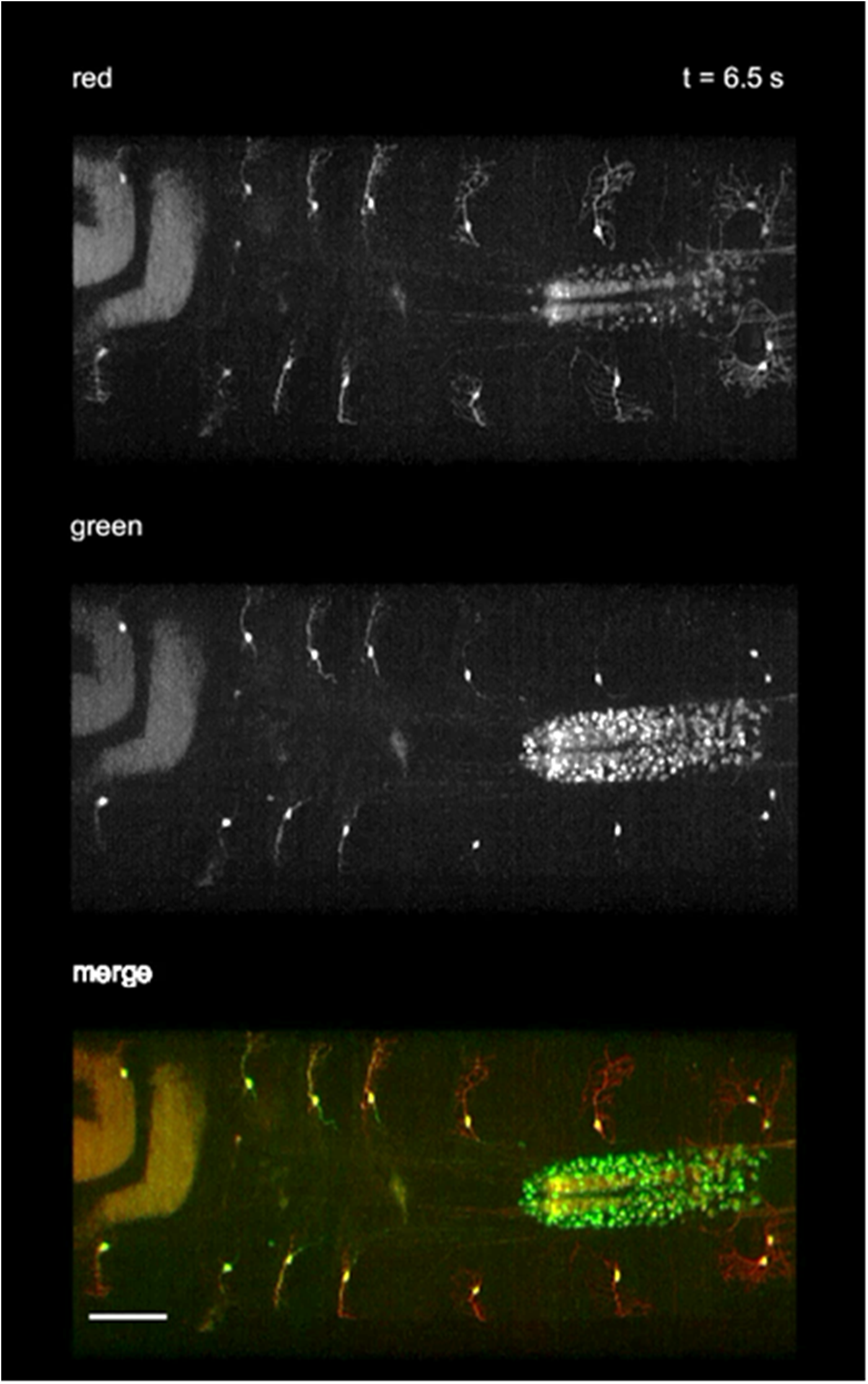
Ventral class I (vpda) GCaMP dynamics during crawling. Vpda neurons are labeled with *410*-*Gal4*, *20XUAS*-*IVS*-*GCaMP6f* (x2), *UAS*-*CD4*-*tdTomato*. Ventral view MIP over a 85 μm depth range of a 153 μm deep volume to exclude gut autofluorescence (square root grayscale). Top panel is tdTomato fluorescence, middle panel is green GCaMP6f fluorescence and the bottom panel is the channel merge. Posterior is to the left. Scale bar=100μm.

**Supplemental movie 4.**
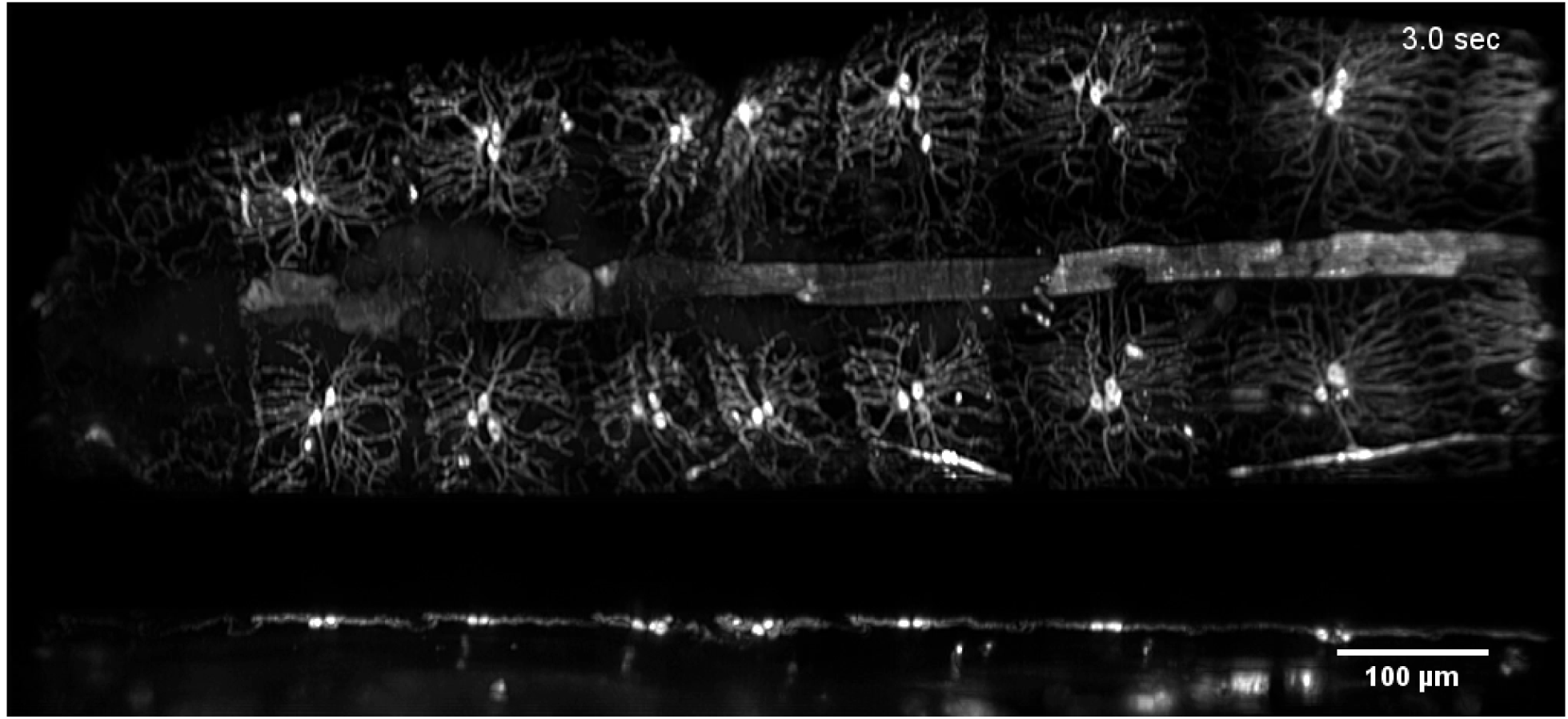
Dorsal class I (ddaE, ddaD) dendrite dynamics during crawling, GFP only. Class I dendrites (and class IV weakly) are labeled by *221*-*Gal4*, *UAS*-*mCD8::GFP*. Top is dorsal view MIP over a 95 μm depth range from a 160μm deep volume to exclude gut autofluorescence, bottom is orthogonal view of dorsal cI neurons ddaD and ddaE (square root grayscale). Posterior is to the left. Scale bar=100μm.

**Supplemental movie 5.**
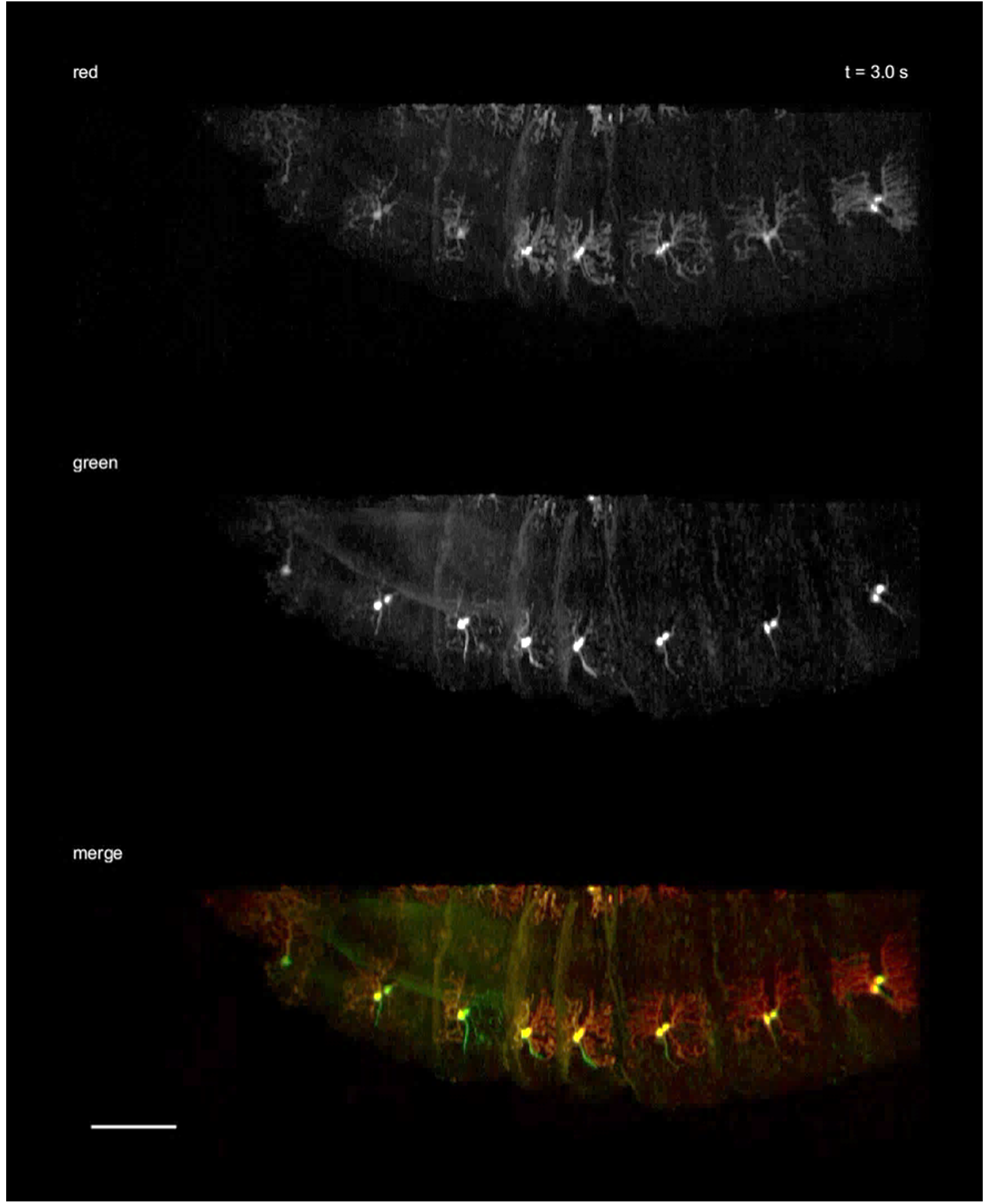
Dorsal class I (ddaE, ddaD) GCaMP dynamics during crawling. cI neurons are labeled with *410*-*Gal4*, *20XUAS*-*IVS*-*GCaMP6f* (x2), *UAS*-*CD4*-*tdTomato*. Dorsal view MIP over a 35 μm depth range of a 200μm deep volume to exclude gut autofluorescence (square root grayscale). Top panel is tdTomato fluorescence, middle panel is GCaMP6f fluorescence and the bottom panel is the channel merge. Posterior is to the left. Scale bar=100μm.

**Supplemental movie 6.**
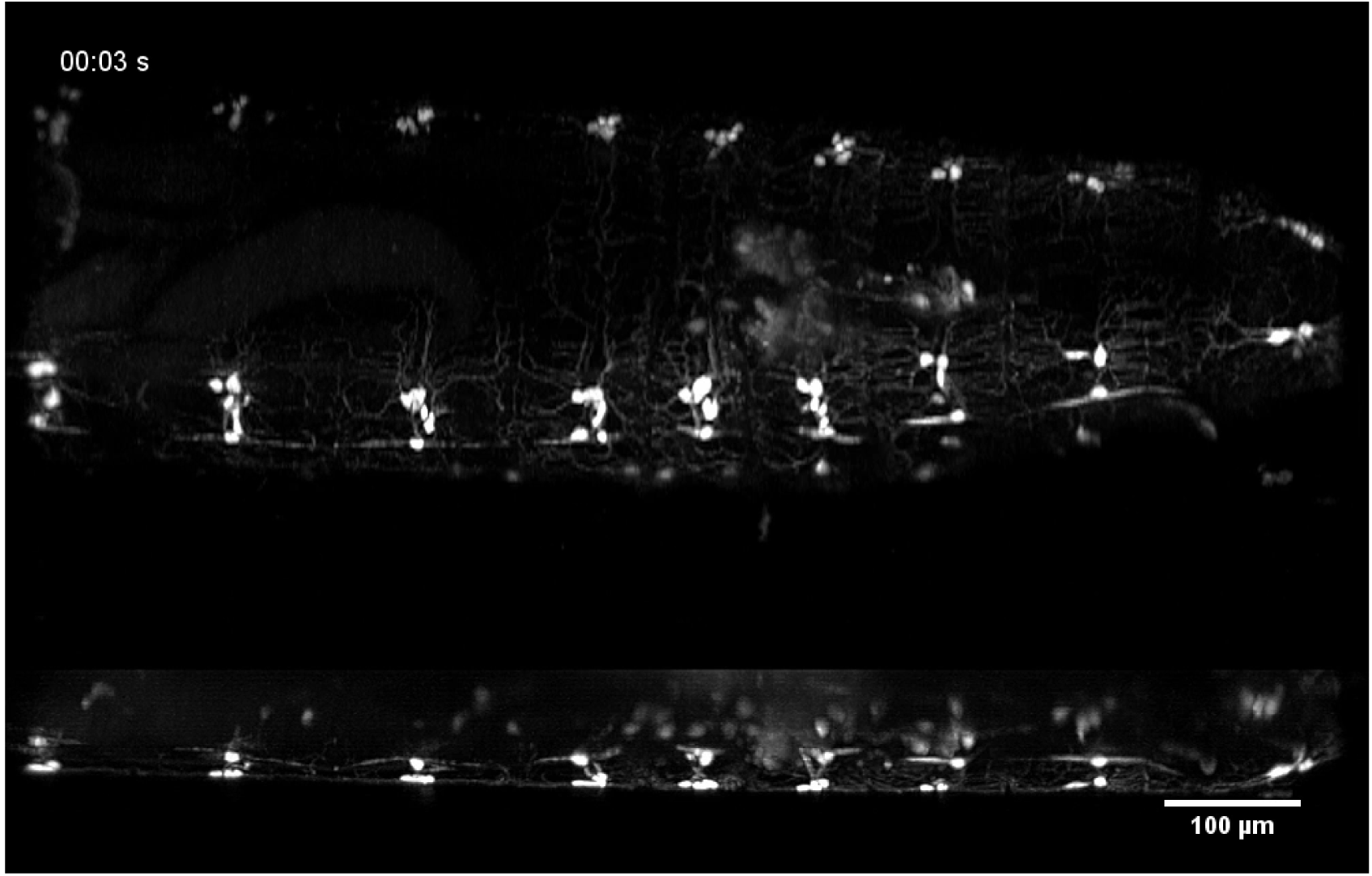
Dorsal cluster dendrite dynamics during crawling, GFP only. dbd, dmd1, ddaE, and ddaD neurons are labeled by *10D05*-*Gal4*, *20XUAS*-*mCD8::GFP*. Top is dorsal view (x-y) MIP over a 90μm depth range from a 165 μm deep volume to exclude gut autofluorescence, bottom is side view (y-z), (square root grayscale). Posterior is to the left. Scale bar=100μm.

**Supplemental movie 7.**
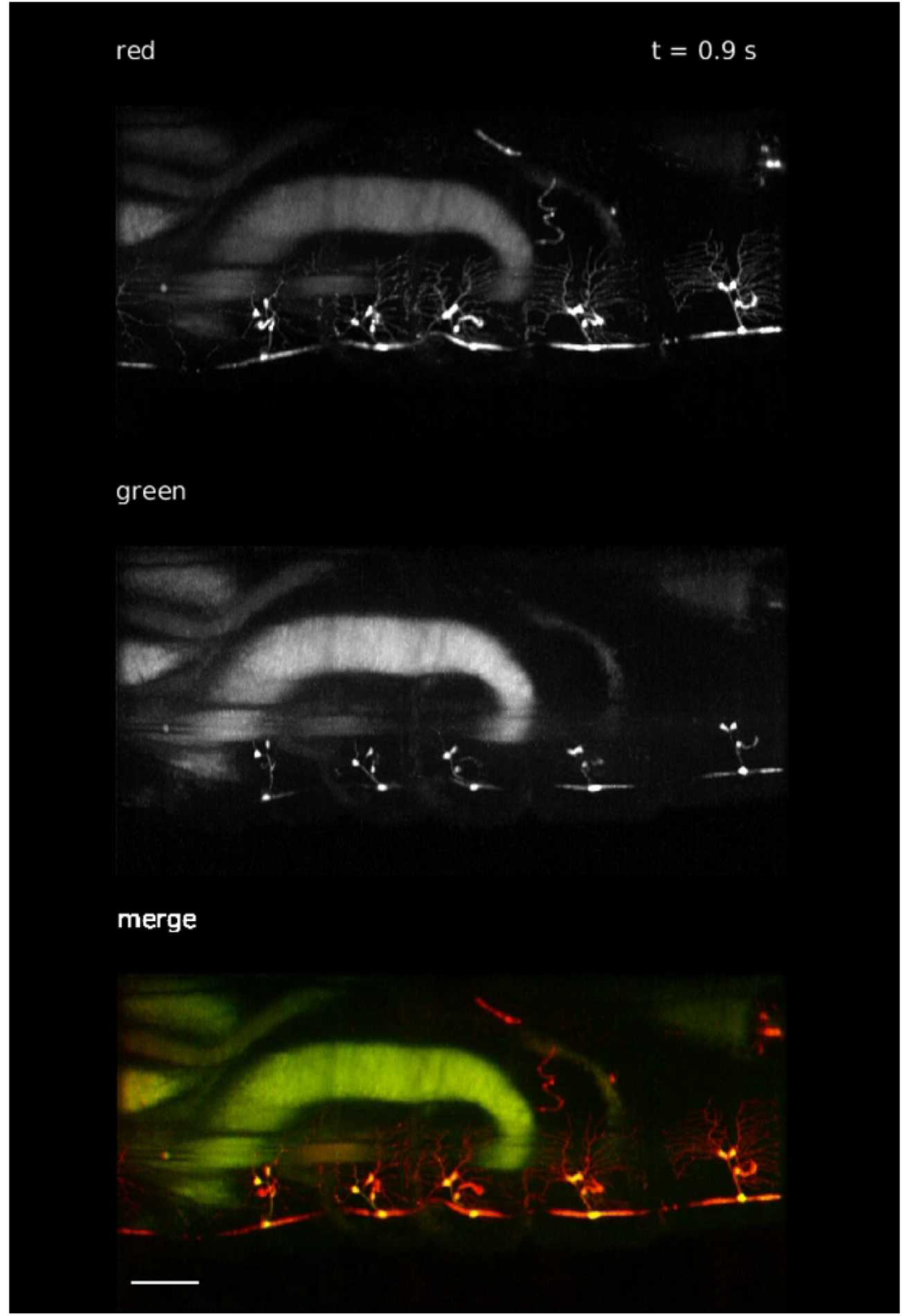
Dorsal cluster GCaMP dynamics during crawling. Dorsal cluster proprioceptors are labeled with *10D05*-*Gal4*, *20XUAS*-*IVS*-*GCaMP6f* (x2), *UAS*-*CD4*-*tdTomato*. Dorsal view MIP over a 100μm depth range of a 200μm deep volume to exclude gut autofluorescence (square root grayscale). Top panel is tdTomato fluorescence, middle panel is green GCaMP6f fluorescence and the bottom panel is the channel merge. Posterior is to the left. Scale bar=100μm.

**Supplemental movie 8.**
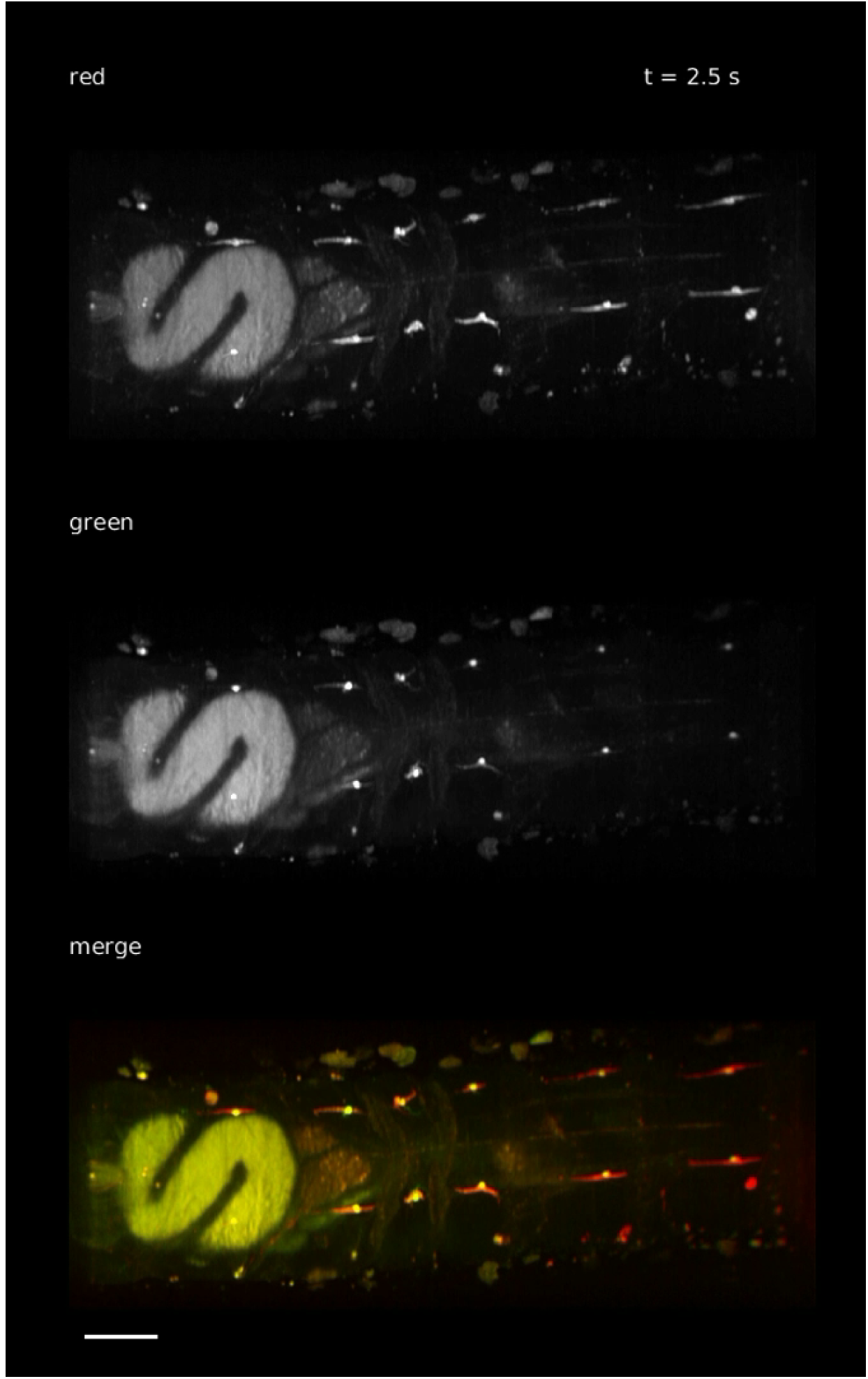
vbd GCaMP dynamics during crawling. vbd neurons are labeled by *1129*-*Gal4*, *UAS*-*CD4*-*tdTomato*, *20XUAS*-*IVS*-*GCaMP6f* (x2). Ventral view MIP over an 80μm depth range of a 160μm deep volume to exclude gut autofluorescence (square root grayscale). Top panel is tdTomato fluorescence, middle panel is GCaMP6f fluorescence and the bottom panel is the channel merge. Posterior is to the left. Scale bar=100μm.

**Supplemental movie 9.**
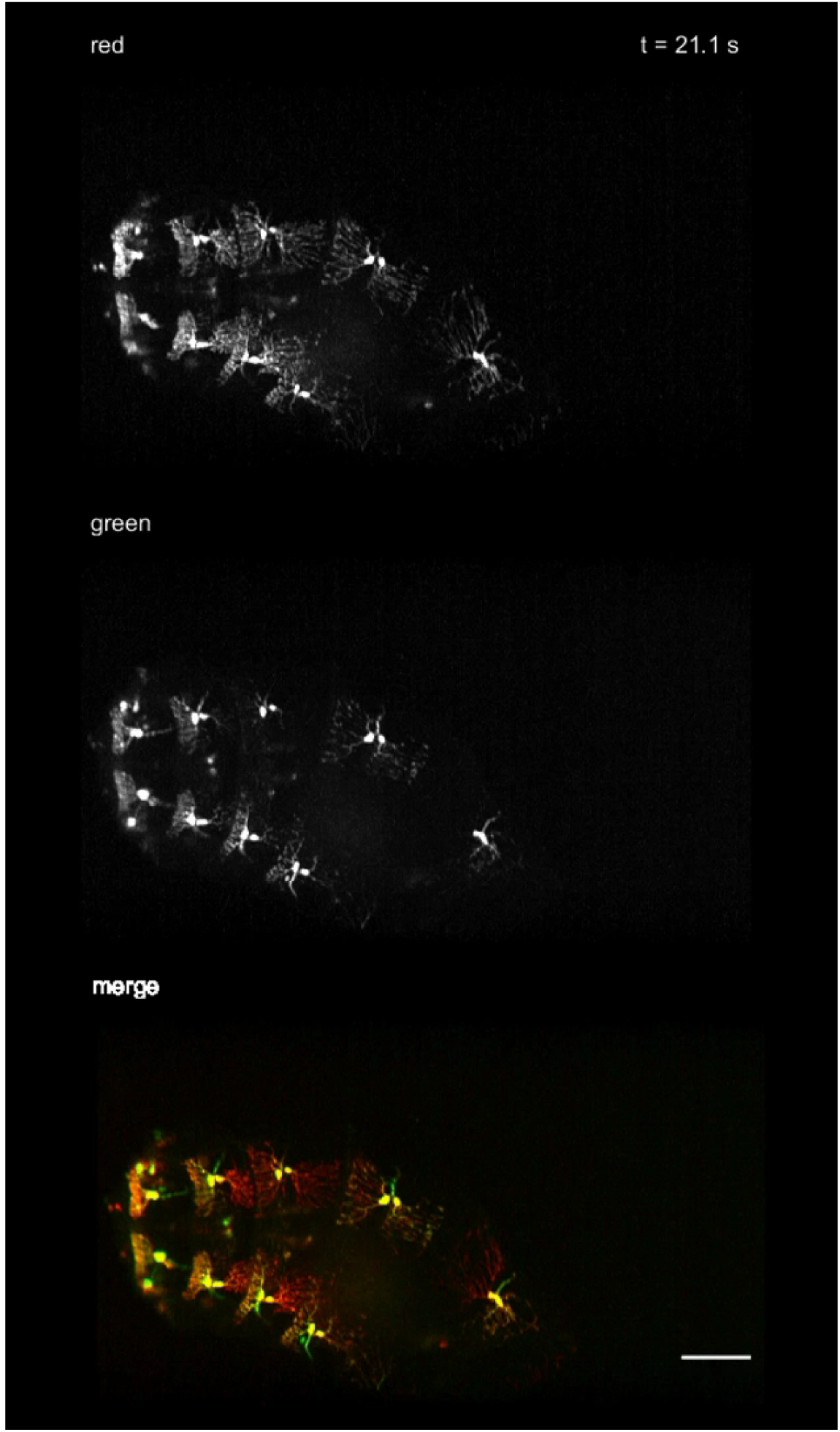
Dorsal class I GCaMP dynamics during head exploration behavior. cI neurons are labeled with *410*-*Gal4*, *20XUAS*-*IVS*-*GCaMP6f* (x2), *UAS*-*CD4*-*tdTomato*. Dorsal view MIP over an 80μm depth range of a 140 μm deep volume to exclude gut autofluorescence (square root grayscale). Top panel is tdTomato fluorescence, middle panel is GCaMP6f fluorescence and the bottom panel is the channel merge. Posterior is to the left. Scale bars=100μm.

**Supplemental movie 10.**
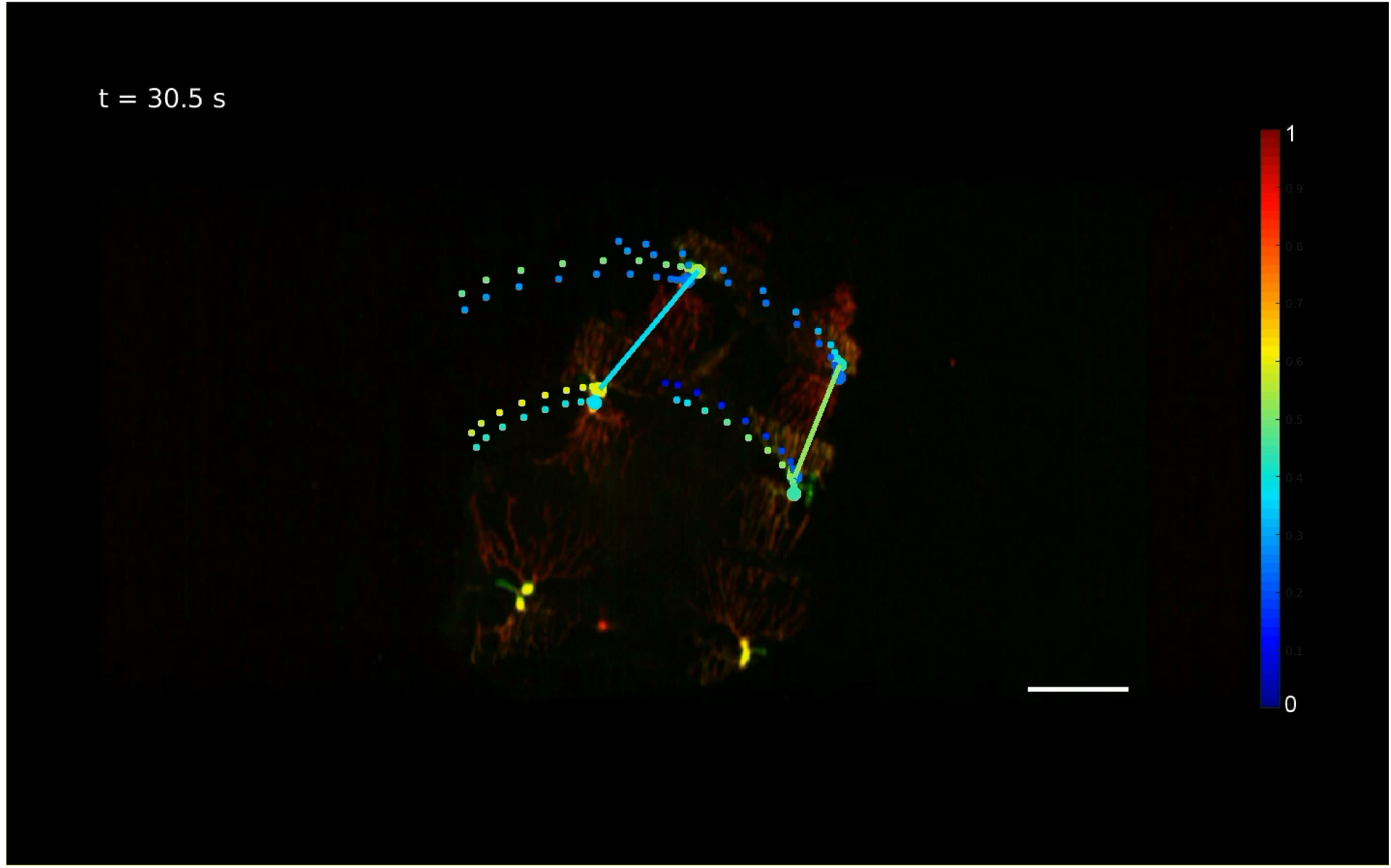
Dorsal class I GCaMP dynamics during head exploration behavior with analysis overlay. cI neurons are labeled with 410-Gal4, 20XUAS-IVS-GCaMP6f (x2), UAS-CD4-tdTomato. Dorsal view MIP over an 80μm depth range of a 140 μm deep volume to exclude gut autofluorescence (square root colorscale). Both ddaD and ddaE neurons from T3 and A1 segment are marked as dots and their GCaMP-tdTomato ratio is color coded to indicate the neuron activity. The distance between two ddaD neurons in each side is also color coded as (red = compression, blue = rest). The trailing spots show prior positions of each neurons and their GCaMP levels. Scale bar=100μm.

